# Temporal genome-wide fitness analysis of *Mycobacterium marinum* during infection reveals genetic requirement for virulence and survival in amoebae and microglial cells

**DOI:** 10.1101/2023.03.22.533734

**Authors:** Louise H. Lefrançois, Jahn Nitschke, Gaël Panis, Julien Prados, Rachel E. Butler, Tom A. Mendum, Nabil Hanna, Graham R. Stewart, Thierry Soldati

## Abstract

Tuberculosis remains the most pervasive infectious disease and the recent emergence of multiple or even fully drug-resistant strains increases the risk and emphasizes the need for more efficient and better drug treatments. A key feature of mycobacteria pathogenesis is the metabolic switch during infection and expression of virulence genes is often adapted to specific infection conditions. This study aims to identify genes that are involved in the establishment and maintenance of the infection. To answer these questions, we have applied Transposon Sequencing (Tn-Seq) in *M. marinum*, an unbiased genome-wide strategy that combines saturation insertional mutagenesis and high throughput sequencing. This approach allowed us to precisely identify the localization and relative abundance of insertions in pools of Tn mutants. The essentiality and fitness cost, in terms of growth advantage and disadvantage of over 10^5^ mutants were quantitatively compared between *in vitro* and different stages of infection in two evolutionary distinct hosts, *D. discoideum* and BV2 microglial cells. We found that 57% of TA sites in the *M. marinum* genome were disrupted and that 568 genes (10.2%) are essential for *M. marinum*, which is comparable to previous Tn-Seq studies on *M. tuberculosis*. The major pathways involved in the survival of *M. marinum* during infection of *D. discoideum* were related to vitamin metabolism, the *esx-1* operon, as well as the mce1 operon.

## INTRODUCTION

*Mycobacterium marinum* is the causative agent of a tuberculosis-like disease in poikilotherms and it is also an opportunistic human pathogen, where the infection is limited to skin and extremities due to growth restriction at about 30°C (Ramakrishnan & Falkow, 1994). In contrast to *Mycobacterium tuberculosis*, which has evolved as a strict human pathogen, *M. marinum* remains a generalist and keeps the capacity to affect a wide range of animal hosts and protozoa. Although *M. marinum* has a larger genome than *M. tuberculosis*, they are phylogenetically (Stinear et al., 2008) related and the processes accompanying infection at a cellular level are similar (Cardenal-Munoz et al., 2018).

Mycobacteria possess a remarkably elaborate cell wall to protect themselves against environmental stresses and chemicals (Brennan, 2003; Jarlier & Nikaido, 1994), and components of the cell wall are well conserved between *M. tuberculosis* and *M. marinum*. The most remarkable is a specific hydrophobic outer layer, called “mycomembrane”, composed of several lipids including mycolic acids (MA), containing between 60 and 90 carbon atoms, and the most abundant lipid, the complex and branched phthiocerol dimycocerosates (PDIMs) (Astarie-Dequeker et al., 2009; Augenstreich et al., 2017). The mycobacterial cell wall is also essential to establish the infection and persist in a non-replicative state during dormancy. The early phase of infection, entry and survival in the macrophage, is determined by key virulence factors, such as the type VII secretion system ESX-1. The ESX-1 system secretes a membranolytic peptide ESAT-6, which acts together with PDIMs (Lerner et al., 2018; Quigley et al., 2017), to produce membrane perforations (J. Smith et al., 2008), leading to the translocation of pathogenic mycobacteria from the Mycobacterium-containing vacuole (MCV) to the cytosol (Houben et al., 2012; Simeone et al., 2012; van der Wel et al., 2007).

Intracellular mycobacteria use host lipids as a carbon and energy source during active growth and dormancy (Barisch & Soldati, 2017). *M. tuberculosis* even has the unique ability to use cholesterol and fatty acids as sole carbon sources *in vivo* and *in vitro* (Nazarova et al., 2019). It has been demonstrated that *M. marinum* is able to use fatty acids derived from triacylglycerols in lipid droplets as well as from membrane phospholipids for its own metabolism and storage (Barisch et al., 2015; Barisch & Soldati, 2017). The metabolic pathways involved in cholesterol and fatty acid catabolism lead to the production of acetyl-CoA, which fuels the TCA cycle, and propionyl-CoA that has to be detoxified and is at least partly used to produce the methyl-branched lipids PDIMs (Lee et al., 2013). To access the host lipids required for its survival, intracellular mycobacteria need specific transporters such as the MCE family transporters that are conserved among many bacteria species including Mycobacteria. Of the four *mce* operons present in the genome of Mycobacteria (*mce1-4*), two have been characterized *in vitro* and *in vivo* as important for uptake and catabolism of cholesterol (*mce4*) and fatty acids (*mce1*) (Wilburn et al., 2018).

The social amoeba, *Dictyostelium discoideum* is an alternative phagocytic host model to study interactions with environmental and pathogenic bacteria (Dunn et al., 2018). The phagocytic pathway is highly conserved and represents a major restriction pathway for bacteria between macrophages and amoebae. Nevertheless, *M. marinum* is able to survive inside *D. discoideum* in a similar way as *M. tuberculosis* in macrophages (Bozzaro et al., 2008; Cardenal-Munoz et al., 2018; Hagedorn & Soldati, 2007). The powerful model *D. discoideum/M. marinum* has been used to demonstrate the role of the ESCRT and autophagy machineries in MCV membrane repair (Lopez-Jimenez et al., 2018), the role of metal poisoning in bacterial growth restriction (Barisch et al., 2018; Hanna et al., 2021) as well as to identify anti-infective compounds (Hanna et al., 2020; Trofimov et al., 2018). Therefore, we use *D. discoideum* and *M. marinum* as a model system to study cell-autonomous defence mechanisms that are relevant to the pathogenesis of tuberculosis (Cardenal-Munoz et al., 2018; Dunn et al., 2018; Tobin & Ramakrishnan, 2008).

The *D. discoideum* host system has been exploited mainly with the use of targeted, candidate-based approaches, to investigate interactions with pathogenic bacteria such as *Salmonella, Mycobacteria, Legionella*, or *Pseudomonas* (Hagedorn & Soldati, 2007; Steinert & Heuner, 2005), but it has also opened to more holistic, data-driven and global approaches to interrogate the host-pathogen interface. Using these approaches, some studies have targeted only the host by proteomic profiling of the MCV (Guého et al., 2019), or both pathogen and host by dual RNA-Seq (Hanna et al., 2019). Similarly, an increasing number of genetic and genomic methods using massive parallel sequencing of transposon insertions (e.g INSeq, HITS, TraDis) have been developed to interrogate the bacterium, including the Tn-Seq method.

Transposon mutagenesis sequencing (Tn-Seq) is a method used to identify essential or relevant genes in bacterial survival or growth under various conditions or selections. The principle is based on identifying the location and the frequency of the transposon insertions in a library by sequencing all the transposon-genome insertion sites. The Mariner transposon, used in the present study, inserts at every TA dinucleotide sequence throughout the genome. Compared to other methods (e. g. TraDis), the depth of sequencing allows the quantification of the relative frequency of individual mutants in the pool before and after various selections (van Opijnen & Camilli, 2010). The Tn-Seq method has been performed with many environmental and pathogenic bacteria (Chao et al., 2013; Gallagher et al., 2011; Kwon et al., 2020; van Opijnen & Camilli, 2013), including *M. tuberculosis* (Barczak et al., 2017; Carey et al., 2018; DeJesus, Gerrick, et al., 2017; Long et al., 2015), before being adapted to other Mycobacteria (Butler et al., 2020; Matern et al., 2020; J. Wang et al., 2014). To date, only one Tn-Seq study performed in *M. marinum* (using the E11 strain) has been published (Weerdenburg et al., 2015). However, comparative genome analysis of *M. marinum* strains isolated from diseased fish or human patients suggests that the E11 strain and the M strain likely belong to two different clusters (Kurokawa et al., 2013) and might have different host ranges. For example, *M. marinum* M is virulent for all the model hosts studied so far, from amoebae to flies, fish and mice (Davis et al., 2002; Talaat et al., 1998). On the contrary, the E11 strain is avirulent for *D. discoideum* (Weerdenburg et al., 2015).

While several screens have been employed to identify genes that are important for mycobacteria survival *in vitro* and *in vivo*, in the present study, we used the reference *M. marinum* strain M, isolated from a human patient, and which is most commonly used as a model for *M. tuberculosis* pathogenesis in fish, flies, macrophages and amoebae (Bozzaro et al., 2008; Cardenal-Munoz et al., 2018). We used the software TRANSIT (DeJesus et al., 2015) to identify essential genes in two infection models, *D. discoideum* and BV2 microglial cells, and compared them to previously described essential genes of *M. marinum* strain E11 and *M. tuberculosis*. We also determined the fitness-impact of genome-wide mutations at various stages of infection in both hosts, which provides insights into the temporal genetic requirements of *M. marinum* to establish an infection.

## RESULTS AND DISCUSSION

### Selection process and distribution of the transposon insertions

The *M. marinum* strain M (hereafter described as *M. marinum*) library was generated with the phage MycoMarT7 vector carrying a *Himar1* transposon that inserts at every TA dinucleotide in the genome, as previously described for *M. bovis* (Butler et al., 2020; Long et al., 2015). For each infection, a frozen aliquot of the *M. marinum* library was grown in 7H9 to generate the reference input sample (hereafter called “Inoculum”) and sequenced. The selection was performed in two consecutive rounds of *D. discoideum* infection. First, to sample the early and late stages of a 48h infection cycle, *M. marinum* were collected from populations of infected *D. discoideum* at 6 hpi, 24 hpi, 48 hpi. Second, in order to further increase the strength of the selection, the 48 hpi sample was grown in 7H9 overnight and used as an inoculum (hereafter called “Inoculum_bis”) for a second round of infection, a sample at 48 hpi was collected (called hereafter “96 hpi”) (Fig 1A).

**Fig 1.**
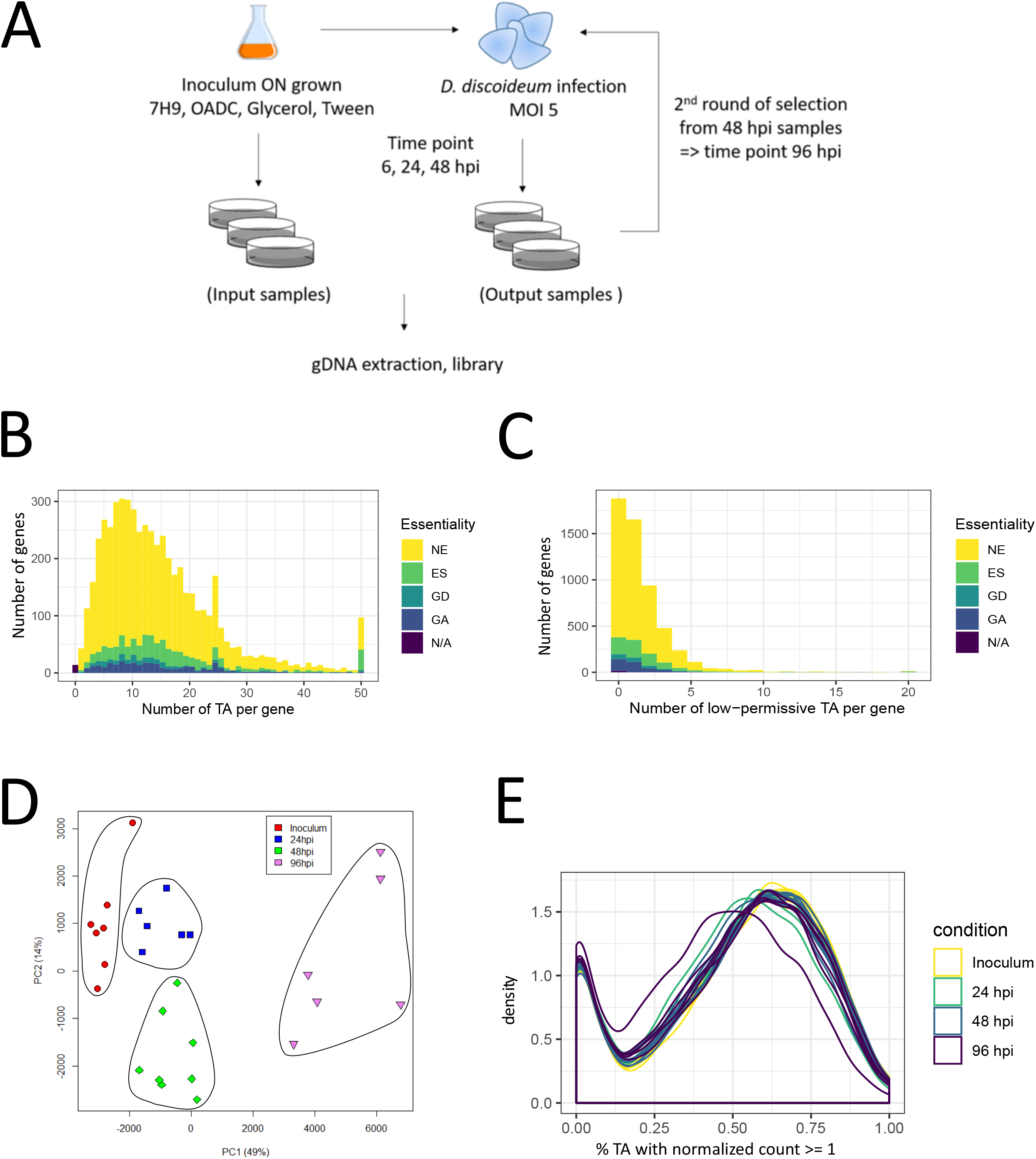
Description of the selection, library coverage and samples selection. **A.** Workflow of *M. marinum* library selection during infection in *D. discoideum. M marinum* stably expressing m-Cherry were thawed and grown overnight. A sample was collected which constituted the “input sample” or inoculum, the remaining culture was used to infect *D. discoideum* AX2 (Ka) and samples were taken over a time-course of 6, 24 and 48 hpi. *M. marinum* were recollected from the 48 hpi samples and used to again infect *D. discoideum*. After another 48 hpi, samples were collected which, for simplicity, are called 96 hpi samples. **B.** Histogram generated from the Hidden Markov Model (HMM) analysis using TRANSIT, representing the essentiality of the 27 samples of *M. marinum* mutant libraries. **C.** Identification of low permissive TA sites in the *M. marinum* genome was performed for each of the 102,057 TA sites. We determined the surrounding +/-3bp sequence and checked whether it matches with the pattern « SGNTANCS » which has been reported to be “low permissive” to transposon insertions. The histogram reports gene essentiality in dependence of the number of low permissive TA sites. **D.** Principal components 1 and 2 of normalized insertion counts, coloured by conditions: Inoculum (n=7), 24 hpi (n=6), 48hpi (n=8), and 96 hpi (n=6). Below the axis label the percentage of variance captured by the respective principal component is denoted.

The genome of *M. marinum* has a length of 6,660,144 bp with one chromosome of 6,636,827 bp and one plasmid of 23,317 bp (Stinear et al., 2008). The genome contains 102,057 TA sites which theoretically represents a possible insertion every 65 bp with 83. 8% (85,478 sites) found in an annotated gene and 16.2% (16,579 sites) in intergenic regions. Based on the sequencing of 7 inoculum samples, we observed an insertion in 57% (58,137 sites) of the TA sites (criteria: average normalised read count ≥1) (Fig 1B, Table S1). Recently, the presence of low permissive sequences with consensus “(GC)GNTANC(GC)” have been highlighted in the *M. tuberculosis* genome (DeJesus, Nambi, et al., 2017). In the *M. marinum* genome, we found 9,370 TA sites (9.2% of the total) embedded in a low permissive sequence. In agreement with that prediction, only 711 (7.6%) of these sites had an insertion while 8,659 (92.4%) had none. We then investigated the impact of these non-permissive TA sites on the gene essentiality status identified by TRANSIT (Fig 1C, Table S2). Note that to retain a maximum of information, a combination of the two available annotations for the genome of *M. marinum* strain M (NC_010612.1 and CP000854.1) was used, resulting in a total number of 5,480 annotated genes. We found that, out of the 651 essential genes identified in the inoculum (Fig S2A), 56 have more than 25% permissive TA-sites, and consequently might be false positives.

The quality control performed on all sequenced samples showed a transposon specificity of 40-86% of the reads generated (Table S1). Principal Component Analysis (PCA) of all 38 validated samples revealed a clustering into two relatively distinct groups between the inoculum and 6 hpi samples, indicative of a potential selection happening during this brief infection time (Fig S1). Because this 6 hours interval is shorter than the generation time of *M. marinum*, two possibilities might explain the difference between the two timepoints. Either the selection occurred upon entry into *D. discoideum* or is due to early killing and digestion of some *M. marinum* mutants. A similar observation was made when comparing the 48 hpi samples to the Inoculum_bis samples used for the second round of infection (Fig S1). This difference might be due to sample manipulation, mainly freezing, thawing and cultivation in 7H9 of the 48hpi samples to prepare the Inoculum_bis, which may have an impact on the proportion of mutants recovered. The 6 hpi and Inoculum_bis samples were removed from further analysis, and the remaining 27 samples were used for the downstream bioinformatic analysis (Inoculum n=7; 24 hpi n=6; 48 hpi n=8; and 96 hpi n=6). A PCA on the normalized read counts per gene was performed for the 27 samples to uncover patterns of similarities among Tn-Seq profiles and to identify clusters of conditions. Altogether, PCA1 (49%) and PCA2 (14%), represent more than half of the variance in the data set. The PCA analysis reveals a clustering of the 27 samples in 4 distinct groups, that match the different infection time points (Fig 1D).

### Identification of the core of essential genes and comparison to other Mycobacteria

The data were analysed using TRANSIT, which is a software that automates the analysis of Himar1 Tn-Seq datasets (DeJesus & Ioerger, 2013). TRANSIT uses a Bayesian method and a Hidden Markov Model (HMM) which is implemented as a 4-state model assigning the regions into 4 categories: (1) essential regions (ES) which are mostly devoid of insertions, (2) non-essential genes (NE) containing read-counts around the global mean, (3) growth-defect genes (GD), and (4) growth-advantage genes (GA) which represent genes with significantly increased or decreased normalised read-counts, respectively.

By definition, essential genes (ES) are genes that do not allow any transposon insertion and therefore, are absent from the original inoculum used to infect *D. discoideum*. Due to biological and technical variations, some mutants might be present under the detection threshold in the inoculum and become significantly represented after selection. As well, some mutants present in the inoculum can disappear during infection because the corresponding genes are essential to grow in *D. discoideum*. Therefore, we defined as “essential core” the genes for which no insertion could be detected in any of the 4 investigated conditions. We identified 651 such genes in the inoculum, 635 genes after 24 h, 630 after 48 h and 704 after 96 h of infection, and 568 genes (10.2%) were common and form the essential core (Fig S2A) (Table S2). Then, we compared the essential core of *M. marinum* to the sets of essential genes previously identified for other mycobacteria species, the *M. marinum* strain E11 (Weerdenburg et al., 2015) and the *M. tuberculosis* H37Rv (DeJesus, Gerrick, et al., 2017) (Fig 2B). First, we identified *M. marinum* orthologs of ES genes reported in the two studies, and then we checked for overlap between these genes and the essential core. This core dataset in our study shared up to 37.3% (N=237) overlap with *M. marinum* E11 (Weerdenburg et al., 2015), and 44.4% (N=311) with the genes classified as ES for *M. tuberculosis* H37Rv (Fig 2B) (DeJesus, Gerrick, et al., 2017). For example, among the common essential genes between *M. marinum* M and *M. marinum* E11, genes encoding proteins involved in DNA replication were identified and more specifically gyrases (gyrB and gyrA), the latter being a known target of anti-TB drugs. On the other hand, genes implicated in nucleotide biosynthesis such as pyrimidine synthase and genes involved in the *de novo* purine synthesis (*purH*) were found to be common essential genes between *M. marinum* M strain and *M. tuberculosis* H37Rv (Table S1). Altogether, 153 genes were common to the “essential core” identified in this study and essential genes both in *M. marinum* E11 and *M. tuberculosis* H37Rv.

**Fig 2.**
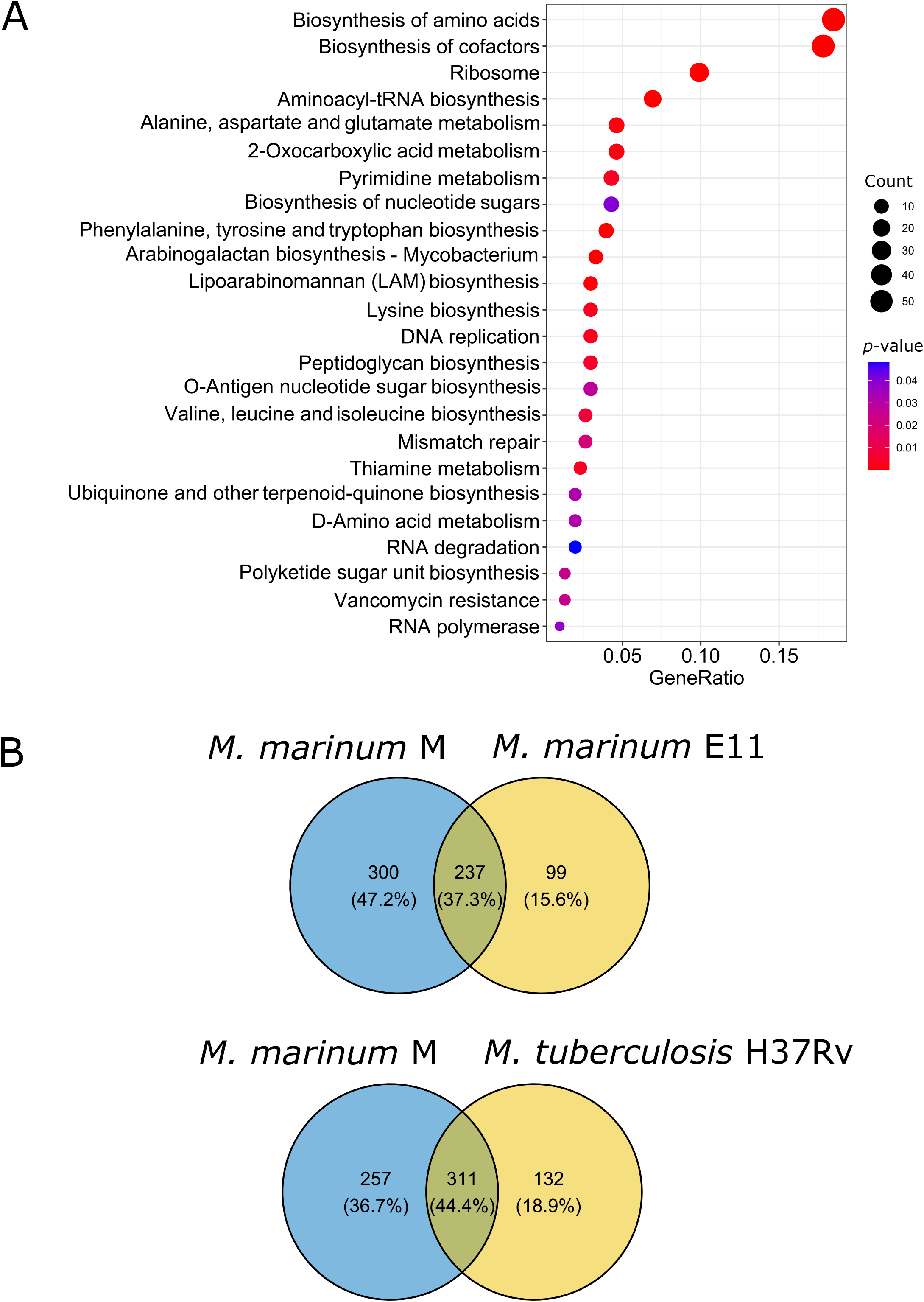
Identification of the “essential core” of *M. marinum* in *D. discoideum*. **A.** KEGG terms enriched in the essential core sorted as a dotplot. Using R and the package clusterProfiler terms were filtered by *p*-value < 0.05. The *p*-value is color coded, and the number of enriched genes found in the respective term are coded by dot size. **B.** Comparison of the “essential core” of *M. marinum* M strain with the *M. marinum* E11 strain and the Mtb H37Rv strain. Comparisons were performed by considering only orthologous genes with a MMAR nomenclature.

The essential core of *M. marinum* was further characterized using a KEGG enrichment analysis to identify functional categories that are overrepresented. KEGG with a p-value < 0.05 were visualized as a dotplot (Fig 2A) and an enrichment map (Fig S2B). The size of the dots represents the number of genes in the KEGG term and the colour of the dots represents the *p*-value (Table S4). The major categories are mostly involved in the intrinsic metabolic pathways essential for the survival of the bacteria *in vitro* and *in vivo*. Among the functional groups identified in this category, DNA replication was significantly enriched as well as genes affecting functions needed for replication fork formation and movement (*dnaB*, DNA primase dnaG encoding DNA primase, and dnaE1 encoding DNA polymerase III), as well as genes implicated in the repair of the lagging strand Okazaki fragments (*polA*, encoding DNA polymerase I, and *ligA*, encoding DNA ligase). Among the identified essential functional groups, genes involved in cell wall synthesis and belonging to the peptidoglycan biosynthesis pathway including *murB* and *ddlA* encoding a D-alanine D-alanine ligase. In addition, the valine, leucine, isoleucine and lysine biosynthesis pathways were enriched including *ilvB1* (MMAR_1720) and *lysA*, respectively. These two genes are known to be essential in *M. tuberculosis* and mutations in these genes are only obtained when the medium is supplemented with the respective amino acids (Awasthy et al., 2009; Pavelka & Jacobs, 1999). Importantly, we identified a set of 42 essential genes specifically at 96 hpi such as *icl1* (MMAR_0792) which encodes for the enzyme isocitrate lyase involved in the TCA cycle. It is known that the disruption of this gene leads to an impaired chronic-phase persistence of *M. tuberculosis* in mice (McKinney JD. et al. 2000). Also, *whiB1* (MMAR_1338), a transcriptional regulator belonging to the WhiB family was identified as essential at this time point. It has been shown that whiB1 plays a role in cell growth and initiation of dormancy in response to nitric oxide (NO) (L. J. Smith et al., 2010). Although, *D. discoideum* does not produce NO to combat bacterial infections, *whiB1* might sense and respond to a more general redox stress response in this infection model. In addition, two genes involved in dihydrofolate biosynthesis (MMAR_5110, MMAR_5111) were identified, the latter being essential for the *de novo* synthesis of folate and the inhibition of its enzymatic activity leads to depletion of the folate pool resulting in growth inhibition and bacterial death (Baca et al., 2000).

### Hierarchical clustering of time-series

To gain global insight into fitness determinants as a function of the stage of infection, we calculated a fold change of the insertions per gene (with at least 5 TA sites and including essential genes) for each time point with the Inoculum as the reference. To these time series, we applied hierarchical clustering and obtained 9 distinct clusters. The profiles of all individually mutated genes were generated and the list of genes is available in the (Table S2). This exploratory analysis allowed us to classify the profiles of all mutated genes in nine different clusters with each cluster containing between 387 (cluster 9) and 681 (cluster 1) genes (Fig 3A). To acquire deeper insights into the biological functions that characterize each cluster, we performed a GO term enrichment analysis on each cluster. Most of the clusters did not show any enriched functional groups with the cut-off values (*p*-value ≤ 0.01) used for this analysis except for clusters 2, 7, and 9 which are represented respectively as enrichment maps (Fig 3 B, C, D). Therefore, we focused on the clusters 2 and 7, which contain mutated genes showing a slight or a drastic decrease in their representation in the pool, respectively, with cluster 7 showing the most pronounced decrease, especially at 96 hpi. On the other hand, cluster 9 shows the opposite trend and includes mutations leading to a continuous increase in fold change over time (Fig. 3A). At first sight, we noticed that the clusters showing decreasing fold changes are mainly related to metabolic processes. Cluster 2 was dominated by pathways implicated in lipid biosynthesis, including triacylglycerol (TAG) and glycerolipid biosynthetic pathways and diacyglycerol O-acyltransferase activity (Fig 3B, Fig S3A). TAGs have mainly been associated with persistence of *M. tuberculosis* and studies showed that they constitute a major source of energy for slow metabolizing populations (Galagan et al., 2013). Among the mutated genes belonging to the latter group, we identified MMAR_1519, the homolog of *tgs1* in *M. tuberculosis*. TAGs accumulate under stress conditions, and a mutation in *tgs1* in *M. tuberculosis* drastically reduced TAG accumulation (Sirakova et al., 2006). In parallel, functional groups involved in various metabolic processes are the most abundant in cluster 7 which shows steeply decreasing representation in the pool over the infection time course. Among these, we identified pathways linked to lipid biosynthesis (*p*hthiocerol *dim*ycocerosates - PDIM) and import of fatty acids (Mce1 operon) both crucial for the survival of *M. tuberculosis* in mice, amino acid metabolism (glutamine), and vitamins biosynthesis (biotin, cobalamin, and folate) indicating an ontogeny-specific metabolic bias for this cluster (Fig 3C, FigS3B). On the other hand, cluster 9 which contains mutations with an increasing representation in the pool, groups genes related to DNA damage binding, acetylation, and aromatic compound catabolic process, which includes genes involved in the shikimate pathway (Fig 3D, FigS3C). Interestingly, clusters 5 showed an alternating trend between GA at 48 hpi and GD at 96 but did show any enriched functional group at the set cut-off value. In order to have a closer look at the functional group enriched in this cluster, we relaxed the set cut-off to *p*-value ≤ 0.05 and performed a GO enrichment analysis. Among the enriched functional groups, we identified DNA-binding transcription factors including MMAR_0408 (*mce1R*) encoding for a putative transcription regulatory protein for the *Mce1* operon.

**Fig 3.**
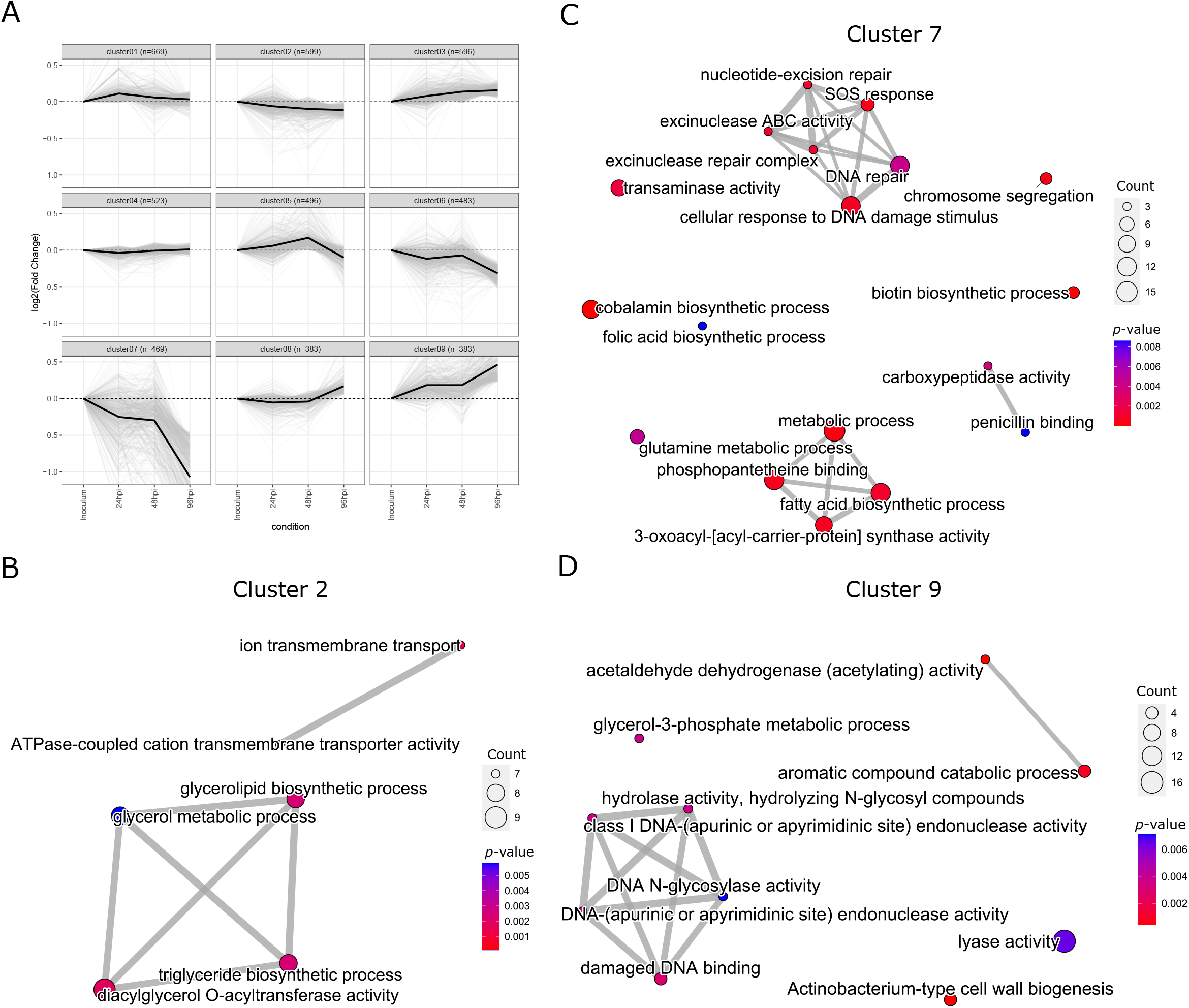
Hierarchical clustering of log2 fold changes in *D. discoideum*. Relative abundance of transposon insertions at various infection timepoints was compared to the initial inoculum. From these binary comparisons, a log_2_fc and a *p*-value were computed. **A.** Genes with at least 5 TA sites and a normalized read count > 1 in any of the samples were selected. The Manhattan distance between each pair of genes across the different timepoints was computed and a hierarchical clustering algorithm with ward.D linkage was run. The resulting clustering tree was cut at a height of 35, and 9 clusters containing between 383 (cluster 8 and 9) and 669 (cluster 1) genes were retained. **B, C, D**. Enriched GO terms in cluster 2, 7 and 9 depicted as an enrichment map. Using R and the package clusterProfiler terms were filtered by *p*-value < 0.01. Edges between nodes represent overlap between the connected terms and the number of enriched genes found in the respective term are coded by dot size.

### Fitness cost of differentially represented mutants during *D. discoideum* infection

The mutations that had impact during infection were identified by comparing mutant abundance (numbers of each unique Tn-Seq read per gene) at the various infection time-points to the inoculum mutant library using the resampling option in the TRANSIT software. Mutants were considered to be enriched or depleted in a condition where the log2 ratio was greater than 0.585 or smaller than −0.585 with a *p-* value ≤ 0.05 (Fig S3E, F, G, Table S3).

Mutations with no impact on fitness in a given condition have fitness scores close to zero. Negative fitness scores (negative log_2_fc) reflect gene disruptions that result in a fitness disadvantage (FD), and positive fitness scores (positive log_2_fc) indicate genes disruptions that result in a fitness advantage (FA). To identify the mutations that have an impact on bacterial growth during infection, we used an HMM resampling method that determines the read counts that are significantly different at 24 hpi, 48 hpi and 96 hpi in comparison to the inoculum (Table S3). Results are presented as volcano plots showing the fold-change (FC) and *p*-values of each pairwise comparison with the inoculum. The overall pattern reveals an unbalanced distribution with more mutations leading to a fitness disadvantage (FD) at 24 hpi (Fig S3E) and 96 hpi (Fig S3F), whereas at 48 hpi (Fig S3G) the effect of mutations is more evenly distributed. Indeed, at 24 hpi, only 70 mutations had a significant impact, all leading to a FD. At 48 hpi, 74 mutations had a significant impact, with 45 leading to a FD and 29 to a FA. At 96 hpi, the highest number of mutations was observed to have a significant impact with 360 conferring a FD and only 15 leading to a FA.

We examined potential relationships among the *M. marinum enriched or depleted mutants* at different timepoints using the STRING database, which curates data from multiple sources including physical as well as functional association between proteins (Legeay et al., 2020). First, *M. marinum* genes were translated to their *M. mycobacterium* orthologs prior to performing the analysis. A colour coding was used to identify the effect of the mutation ranging from dark blue (mutations leading to FD) to dark red (mutations leading to FA). The different timepoints are represented as circles around the gene name where the inner circle corresponds to 24 hpi and the most outer layer corresponds to 96hpi (Fig. 4). This analysis revealed that majority of transposon insertions and their corresponding protein subnetworks confer a clear FD, such as the ESX-1 complex, iron homeostasis, DNA repair, cobalamin and biotin metabolism. Few dispersed genes confer a clear FA, such as relA or Rv3816c.

**Fig 4.**
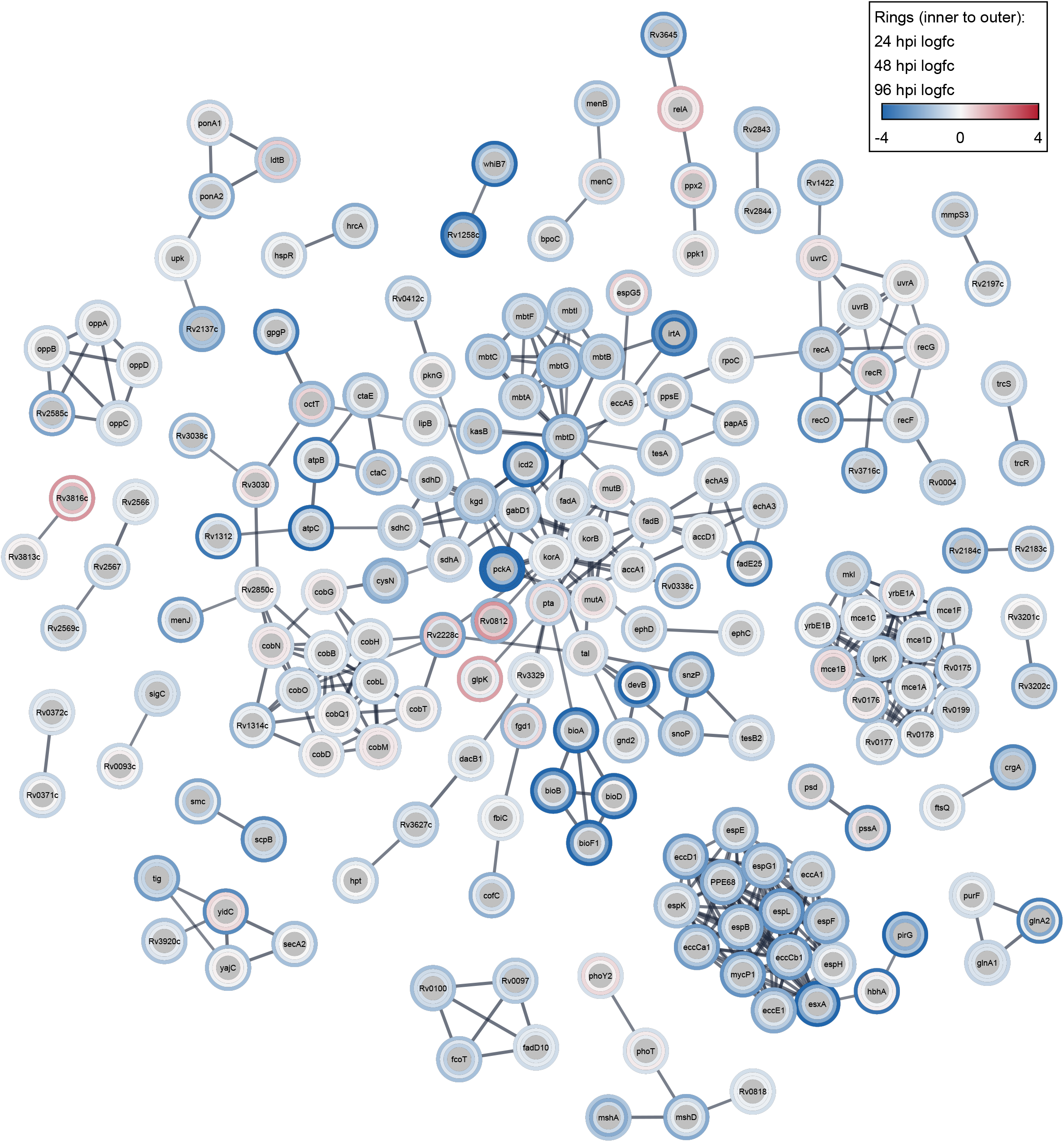
Protein interactions of FA/FD genes over time. The figure displays a protein-protein interaction network of Mtb orthologs of FA or FD genes, retrieved from the STRING database (genes filtered for absolute logfc >= 0.585, p-value <= 0.05, projected on the full STRING network, confidence cut-off of 0.8, singletons were omitted, query organism: *Mycobacterium tuberculosis* H37Rv). A colour coding was used to identify the effect of the mutation ranging from dark blue (mutations leading to FD) to dark red (mutations leading to FA). The color scale in the layered rings represents from inside to outside the logfc of 24, 48 and 96 hpi respectively compared to the Inoculum. Genes thresholded as described in Table 1 were submitted to GO term enrichment using R and the package clusterProfiler. Terms were filtered by *p*-value < 0.01. Edges between nodes represent overlap between the connected terms, whereas the *p*-value is color coded and the number of enriched genes per term is coded by the dot size. **A.** Enriched GO terms for FA and FD at 24 hpi **B.** Enriched GO terms FA and FD at 48 hpi **C.** Enriched GO terms FA at 96 hpi **D.** Enriched GO terms FD at 96 hpi

Interestingly, genes coding for the T7SS subunits were identified which formed a high confidence interactome including MycP1, a protease involved in regulation of ESX-1 secretion and virulence, and EccCb1, a member of the FtsK/SpoIIIE-like ATPse family which provides the energy to transport proteins across the mycobacterial membrane. One important information that could be identified by looking at the STRING analysis is the mutation effect at the different infection timepoints. Although most of the mutations show a linear phenotype, some mutations showed an alternating phenotype such as *glpK* (FD at 24 hpi and 48 hpi, FA at 96 hpi) which has been shown to be involved in a special type of drug resistance called phase variation type (Safi et al., 2019). The opposite phenotype was observed for *uvrC* (FA at 24 hpi and 48 hpi, FD at 96 hpi) a gene involved in nucleotide excision repair pathway (NER) (Fig. 4).

A GO enrichment analysis was performed to unravel specific signatures and determine a temporal trend of the fitness cost of disrupted genes. We submitted significant genes (*p*-value ≤ 0.1, absolute log_2_fc ≥ 0.585) to the enrichment analysis (24 hpi FA and FD combined, Fig S4A; 48 hpi FA and FD combined, Fig S4B; Table S4). Consistent with longer lasting selection the 96 hpi dataset presented the highest number of significant genes. To obtain comprehensive enrichment results, we submitted FA and FD separately to the enrichment analysis, maintaining the *p*-value ≤ 0.1 for FA genes (Fig S4C), but tightening the cut-off to *p*-value ≤ 0.01 for FD genes (Fig S4D, Table S4). The enriched pathways during the different infection timepoints were mainly associated with metabolic processes, cell wall biogenesis, cell cycle, and protein export. The 24 hpi timepoint showed a considerably limited number of enriched pathways including the menaquinone pathway which is part of the electron transport chain and the cell division pathway with genes involved in chromosome structure and partitioning. Interestingly, some functional groups emerged at late time points (48 and 96 hpi) which coincides with *M. marinum* proliferation in the cytosol of its host. Among the most enriched pathways and common to these two time points, we identified the vitamin B6 biosynthesis pathway, and the biotin biosynthesis pathway which is required to support the biosynthesis of methionine, glycine, serine, pantothenate, purines, and thymidine (Minato et al., 2015). Finally, at 96 hpi with the highest number of differentially transposon mutated genes, among the functional group identified in the FD category, a cluster of highly overlapping nodes related to the cellular response to stress (SOS and DNA damage), suggesting that these processes may be integrated during *D. discoideum* infection. Among these, the DNA repair pathway, many helicases (RecG, MMAR_1358) and the well-studied *recA*, a gene involved in regulation of nucleotide excision repair and in induction of the SOS response, are present (Müller et al., 2018). Mycobacteria that reside in host macrophages, are exposed to frequent DNA damaging assaults by a variety of endogenous and exogenous factors. Consequently, activation of DNA repair systems is necessary to ensure an efficient error-free transmission of genetic material. Interestingly, the fatty acid biosynthesis pathway and the related pathways as shown on the enrichment map were also identified at 96 hpi in both FA and FD categories (Fig 4C, D). First, the enrichment of these terms clearly demonstrates that fatty acid metabolism in *mycobacteria* is vital to the survival of the bacterium in the host, which is consistent with the waxy mycomembrane being a key feature of its virulence. Additionally, the presence of processes linked to fatty acid biosynthesis GO terms enriched in both FA and FD categories indicates that the requirements of these metabolic pathways is highly finetuned during infection as opposed to growth *in vitro*. Among the genes that were found in these functional groups, we identified the polyketide synthase pks15/1 which is involved in PGL synthesis.

### The ESX-1 operon and genes implicated in vitamin and lipid metabolism are required for *M. marinum* survival in *D. discoideum*

The overall pattern shows that the majority of the insertions had no impact and are considered as neutral (NE), whereas only few insertions led to a FA or FD in intracellular growth (Fig 5A, B, C) (Table S2). In order to monitor and visualise the even distribution of chromosomal insertions in the tested conditions, and identify potential “hot spots” of mutations and potentially important operons, we plotted the log_2_fc of all genes along their genome position and colour coded the genes by *p*-value (*p*-value ≤ 0.1 or *p*-value ≤ 0.01), at 24 hpi (Fig 5A), 48 hpi (Fig 5B) and 96 hpi (Fig 5C) (table S3). At 24 hpi, most of the mutations were found in genes coding for conserved or hypothetical protein. The gene *pckA* (MMAR 0451) involved in the tricarboxylic acid cycle (TCA) and the *esx-1* operon (10 genes between MMAR_5440 to MMAR 5449) involved in MCV escape, standing out with a relatively low, negative log_2_fc and high significance (p-value 0-0.01). At 48 hpi,*pckA* and the esx-1 operon (with 10 genes affected) with a decreasing negative log_2_fc. In addition, we found mutation in genes involved in lipid metabolism such as the isocitrate lyase *icl2* (MMAR_0158) which plays a pivotal role in regulating carbon flux between the tricarboxylic acid (TCA) cycle, the glyoxylate shunt and the methylcitrate cycle at high lipid concentrations (Bhusal et al., 2019). In addition, we identified *cpsA* (MMAR_4966), a LytR-CpsA-Psr (LCP) domain-containing protein which is required for *M. tuberculosis* to evade killing by NADPH oxidase and LC3-associated phagocytosis (Köster et al., 2017). At this time point, some mutations confer a FA for instance, the ABC transporter encoded by *pabC* (MMAR_4873), as well as *proC* (MMAR_0826) or *fbiB* (MMAR_1280) whose functions are not fully elucidated. At 96 hpi, which reflects the completion of approximately two infection cycles and therefore includes bacterial escape to the cytosol and cell-to-cell dissemination, we found mutations in some previously mentioned genes with a higher log_2_fc than at 48hpi, for *icl2, pckA* and the *esx-1* operon (with mutations in 16 genes of the operon becoming significant at 96 hpi compared to 10 genes at 48 hpi). Additionally, we found some well-known virulence factors involved at different steps of the mycobacterial infection. A mutation in the gene of the heparin-binding hemagglutinin (MMAR_0800: *hbha*) involved in the adhesion of bacteria to host cells; mutations in a gene of the Type 7 Secretion System (T7SS) ESX-5 (MMAR_2680) involved in transport of cell envelope proteins; mutations in different genes implicated in lipid metabolism (MMAR_1778: *tesA*, MMAR_2124: *relA*, MMAR_4966: *cpsA*, MMAR_0258: *fadD10*, MMAR_0319: *fadD7*, MMAR_4864: *mshD);* mutation in the *whiB7* transcriptional regulatory factor (MMAR_1365) or in a gene encoding a serine/threonine protein kinase which is involved in *M. tuberculosis* viability *in vitro* and in infection models (MMAR_0713: *pknG*) (Cowley et al., 2004). More interestingly, at 96 hpi, mutations in genes of the *mce1* operon (MMAR_0410 to MMAR_0417), a putative fatty acids transporter, appeared to induce a relatively strong FD. In addition, mutations in genes involved in vitamin metabolism e.g. biotin (MMAR_2383 to MMAR_2387) or cobalamin (MMAR_1883 to MMAR_1885) also induced a strong FD. Regarding the few mutations that lead to a FA, we found a nitrate response regulator (MMAR_4782: *NarL*), a glycerol kinase (MMAR_5208: *glpK*), an acetyltransferase (MMAR_5379) and some hypothetical secreted or regulatory proteins (MMAR_1939, MMAR_1165) (Table S3).

**Fig 5.**
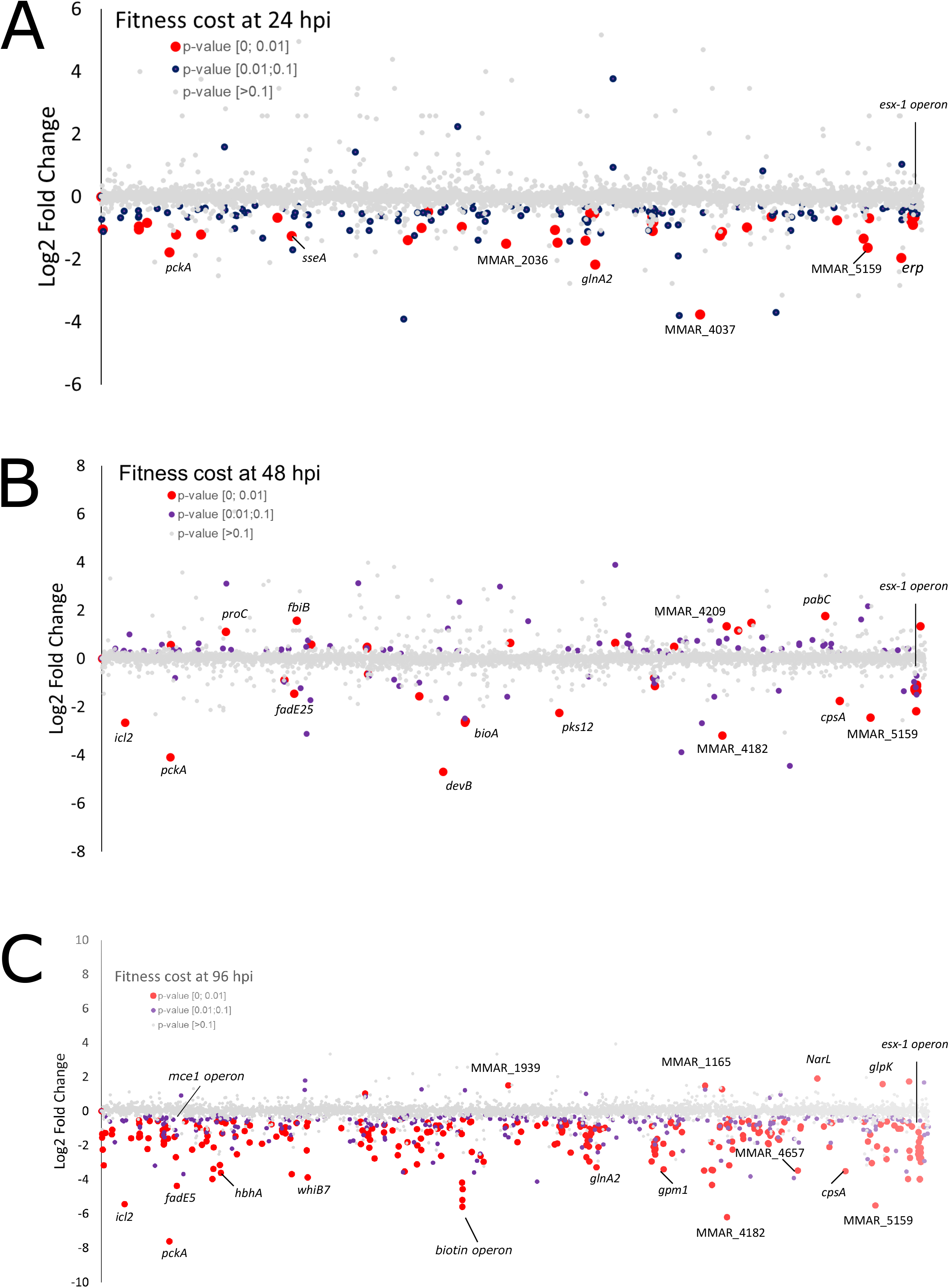
Fitness cost of *M. marinum* genes in *D. discoideum* over time and with respect to genome position. The log_2_fc was obtained by comparing the time points 24hpi, 48hpi and 96 hpi to the inoculum. Genes are represented by dots which are color coded according to the *p*-value associated to the log_2_fc. Red: <0.01, purple: between 0.01 and 0.1, grey: > 0.1. Plotting the log_2_fc in dependence of genomic position (x-axis) reveals the fitness impact of operons in each condition: **A.** 24 hpi, **B.** 48 hpi, **C.** 96 hpi. Striking loci and operons are highlighted by labels and circles, such as the ESX-1 and mce1 operons or genes associated with biotin metabolism.

### Fitness cost of mutations in the *mce1* and *mce4* operons during *D. discoideum* infection

In bacteria, functionally related genes are often organized in operons that ensure the coordinated expression of all the constituent genes and the stoichiometry of the respective proteins/enzymes. As described above, we found pathways linked to lipid biosynthetic processes in both FA and FD enriched GO terms (Fig S4C, D). This prompted us to see whether several operons as organisational units show similar trends for fitness cost (FA or FD). We looked at the impact of mutations in the canonical mycobacteria virulence locus *esx-1* (Fig 6A), which contributes to disruption of the macrophage and amoebae MCV membrane upon infection (Augenstreich et al., 2017; Conrad et al., 2017; Hagedorn et al., 2009), the two operons *mce1* and *mce4* (Fig 6B), and PDIM and PGL biosynthesis genes (Fig 6C). The analysis of the *esx-1* operon and associated genes showed that most of the mutations led to FD, confirming and that this secretion system is required for the growth of *M. marinum* during infection. It is notable that the FD conferred by most of mutations increases during the infection course reaching 16-fold difference for genes encoding EsxA (ESAT-6) and EsxB (CFP10) at 96 hpi (Fig 6A, table S3).

**Fig 6.**
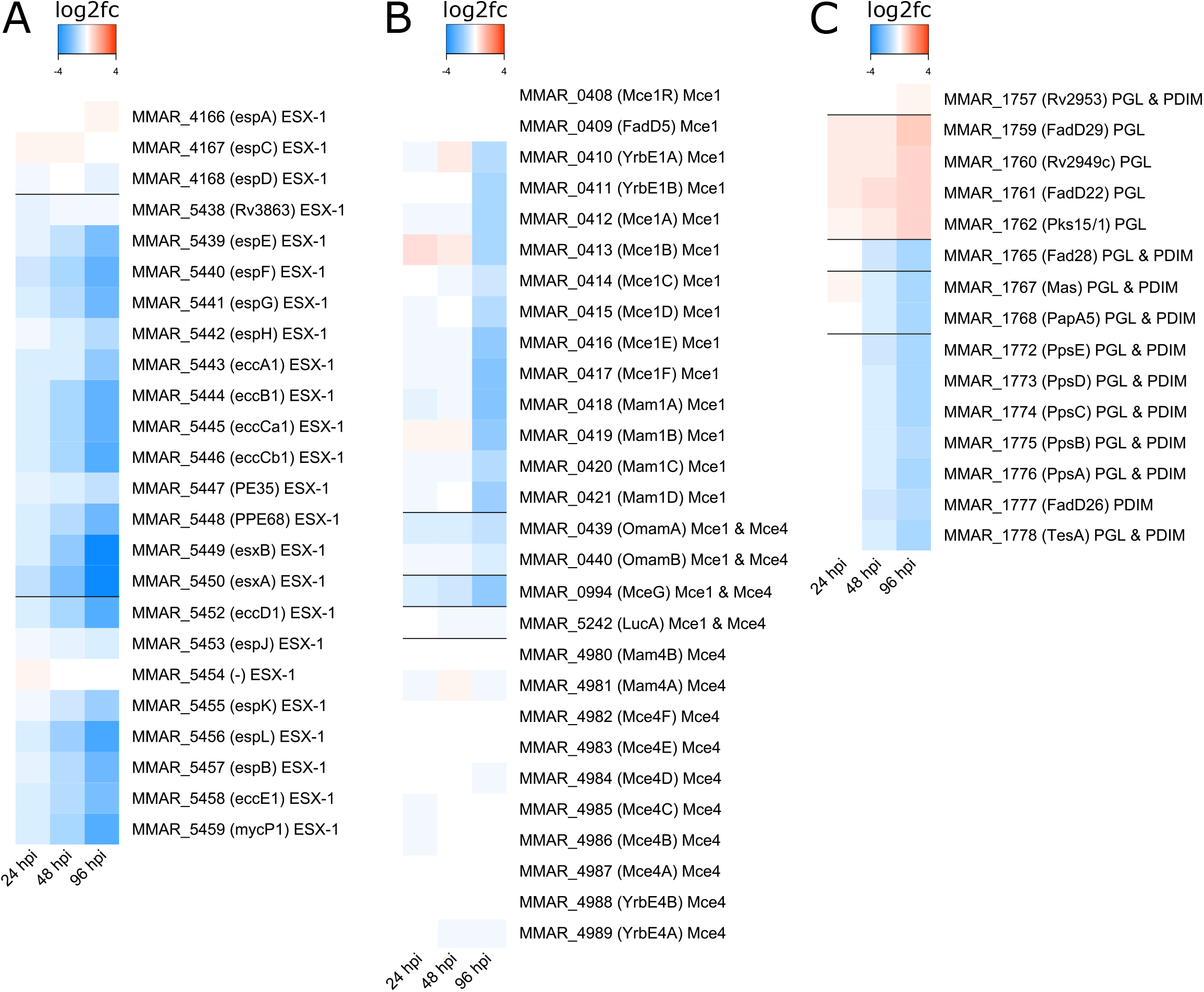
Fitness cost over time for selected genes and operons of *M. marinum* in *D. discoideum*. The color coded log_2_fc in dependence of conditions reveals fitness costs within operons. Operons and associated genes were selected. **A:** *esx-1* operon system, **B:** *mce1* and *mce4* operons systems, **C:** Genes associated with Phthiocerol dimycocerosates (PDIM) and phenolic glycolipids (PGL) synthesis. The labels include the *M. marinum* MMAR nomenclature, in brackets the Mtb orthologue (as indicated on mycobrowser.epfl.ch, with indicating absence of an Mtb orthologue), and the associated function.

During infection, mycobacteria survival, including central metabolism and cell-wall biosynthesis, necessitates lipid import from the host *via* Mce transporters (F. Zhang & Xie, 2011). The heatmaps showing the fitness trends of the *mce1* operon, encoding a fatty-acid transporter, and of the *mce4* operon, encoding a (chole)sterol transporter, are represented on Fig 6B. Both are composed of two putative permeases (*mce1:* MMAR_0410, MMAR_0411 – *mce4:* MMAR_4989, MMAR_4988) and six cell-wall associated proteins (*mce1*: MMAR_0412 to MMAR_0417 - *mce4*: MMAR_4987 to MMAR_4982). In addition, the role of accessory proteins in the stabilization and function of both Mce transporters have been described, including *LucA* (MMAR_5242/Rv3723), Mce-associated membrane proteins (MAMs) and an orphaned MAM (OMAM), as well as *mceG* (MMAR_0439/Rv0199), encoding a putative ATPase (Nazarova et al., 2017, 2019; Perkowski et al., 2016) (Fig 6B). Note that, only the *mce1* operon has a regulator encoded by *mce1R* (MMAR_0408). An inspection of the data revealed that all the mutations in genes belonging to the *mce1* led to FD starting at 48 hpi and reaching the highest FD fold change at 96hpi, except for Mce1R (MMAR_0408) and Fad5 (MMAR_0409) which showed a neutral phenotype. Most importantly, mutations in the gene encoding the ATPase MceG that is predicted to interact with all the Mce transporters, and to be the only cytoplasmic protein, led to a FD phenotype (Rank et al., 2021). These results suggest that mutations in the *mce1* operon and accessory genes mainly impact the growth and survival of *M. marinum* at late times of infection (Fig 6B). However, mutations in the *mce 4* operon showed irregular phenotypes with some mutations leading to a neutral effect or to a slight FD. This finding is surprising, as mutations in the *mce4* operon, previously described as essential for *M. tuberculosis* to survive in the host, seem not to profoundly impact the bacterium during infection in *D. discoideum*. It is noteworthy, that mutations in two genes encoding OMAMs, *omamA* and *omamB*, exerted a FD at 96 hpi (Fig 6B). This result is consistent with data from genetic analyses showing that OmamA and OmamB are required for cholesterol import, despite the lack of impact of most mutations in the *mce4* operon (Rank et al., 2021).

The presence of PDIMs in the mycobacteria envelope has proven to be essential for several key steps of the infectious cycle. More precisely, this component plays a pivotal role in bacterial entry into the macrophage, control of phagosome acidification, phagosomal escape, and use of host cholesterol as a carbon source (Astarie-Dequeker et al., 2009; Osman et al., 2020). On the other hand, Phenolic glycolipids (PGLs) are polyketide-derived virulence factors that are able to limit the capacity of activated macrophages to induce nitric oxide synthase (iNOS) and generate NO upon mycobacterial infection (Oldenburg et al., 2018). In contrast to the *mce1* and *mce4* operons, almost all the mutations in genes belonging to either PGL or PDIM biosynthesis pathway show a continuous increase or decrease of fitness (Fig 6C). This empirically supports the notion of both PGL and PDIM playing a role already early during the infection. The synthesis routes of both PDIM and PGL are connected. A closer look into the mutation in genes belonging to PDIM and PGL pathway led to a FD for PDIM, whereas mutations in genes specific to the biosynthesis of PGL (including the operon MMAR_1759 to MMAR_1762) led to a FA phenotype. This counterintuitive positive impact of loss of PGL synthesis might be due to substrate rerouting to the PDIM pathway upon interruption of the PGL pathway, increasing total PDIM synthesis, boosting virulence and thus effectively leading to FA of these transposon mutants.

### Comparative Tn-Seq shows common fitness impact and essentiality in amoebae and murine microglial host cells during *M. marinum* infection

In order to confirm that *M. marinum* uses common molecular mechanisms and virulence factors to survive in *D. discoideum* and in mammalian phagocytes, we applied a similar selection to the *M. marinum* mutant pools by infecting BV2 murine microglial cells. Based on prior knowledge about the infection cycle in the BV2 microglial cells (Barisch et al., 2015; Hagedorn et al., 2009; Trofimov et al., 2018), a single time point at 48 hpi was collected before cells died from the productive infection. In total, seven samples were used for the analysis, four belonging to the inoculum and three at 48hpi. As for *D. discoideum* experiments, a quality control of reads was performed, showing a transposon specificity of 58-82% (Table S2). Then, a PCA was computed and plotted, showing a distinct clustering of the inoculum and 48 hpi samples, with PC1 and PC2 accounting for more than 75% of the variation in the entire data set (Fig S4A).

To identify mutations that have an impact on intracellular bacterial growth during BV2 infection, we used TRANSIT to compare read counts at 48 hpi to the inoculum, analogously to the procedure applied in experiments with *D. discoideum*. Interestingly, the number and frequency of mutations represented in the pool after BV2 infection is much lower in comparison to the pool after *D. discoideum* infection at 96 hpi. Inspection of all genes on a volcano plot (Fig 7A) clearly shows that the number of mutations leading to FD is higher compared to the mutations leading to FA, which is consistent with the results obtained for the *D. discoideum* infection at 96 hpi (Fig 7A). Markedly, the mutation with the highest FD score (32-fold change) in comparison to the inoculum was in a gene coding for hydantoin racemase (MMAR_4457), whereas the mutation with the highest FA score (10-fold increase) was the aspartate decarboxylase *PanD* (MMAR_5104), which is required for the biosynthesis of the essential cofactor coenzyme A and the target of the *M. tuberculosis* first line antibiotic pyrazinamide (Fig 7A). Next, a gene ontology enrichment analysis was performed on differentially represented mutations at 48 hpi, using *p*-value ≤ 0.1 and absolute log_2_fc ≥ 0.585, enriched terms were filtered by *p*-value < 0.01 (Fig 7B, C). Similar functional groups were enriched during BV2 infection compared to *D. discoideum*, and these were mainly related to vitamin biosynthetic processes such as biotin and cobalamin (Table S4). In addition, an rRNA methylation cluster was identified in the enrichment map, which hints towards a possible post-transcriptional modification involvement during infection (Fig 7B, C). On the other hand, some functional groups that were enriched after the selection in amoebae were not highlighted here, including TCA cycle and lipid catabolic processes (Table S4).

**Fig 7.**
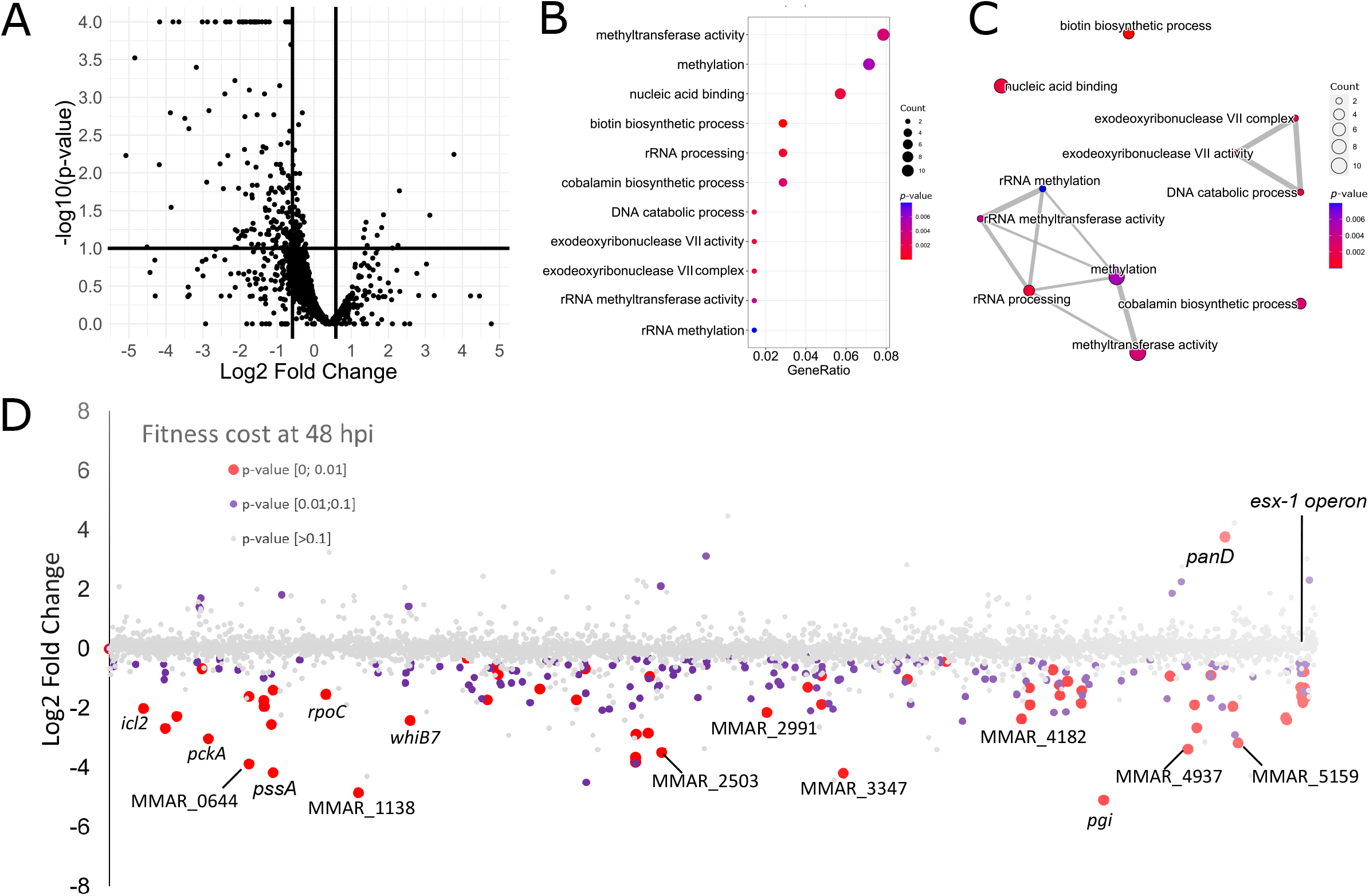
Fitness cost over time of *M. marinum* in BV-2 microglial cells. **A.** Volcano plot of 48 hpi versus the inoculum. The x-axis shows the log_2_fc and the y-axis the -log^10^ of the associated *p*-values. Thresholds used to filter genes for enrichment analysis are depicted as thick black lines: log_2_fc ≥ 0.585 and ≤ 0.585 (corresponding to 1.5fold change) and *p*-value ≤ 0.1. For visualization *p*-values < 0.0001 were capped to 0.0001. **B, C.** Thresholded genes at 48 hpi were submitted to enrichment analysis using R and the package clusterProfiler. Terms were filtered by *p*-value < 0.01. In C, edges between nodes represent overlap between the connected terms, enriched genes found in the respective terms are coded by dot size. **D.** Fitness cost at 48 hpi in BV-2 microglial cells and with respect to genome position. Genes are represented by dots which are color coded according to the *p*-value associated to the log_2_fc. Red: <0.01, purple: between 0.01 and 0.1, grey: > 0.1. Plotting the log_2_fc in dependence of genomic position (x-axis) reveals the fitness impact of operons at 48 hpi.

In order to monitor and visualise the even distribution of chromosomal insertions in the tested condition. Analogously to Fig 5, the log_2_fc was plotted along the genomic position of the respective gene and colour coded by *p*-value ≤ 0.1 or *p*-value ≤ 0.01. The mutations’ fitness cost showed a similar trend as in *D. discoideum* infection in which most of the mutations led to a FD and only one mutation in the *panD* gene (MMAR_5104) led to a significant FA. Among the mutations leading to FD, we found *icl2, pckA, whiB7*, genes involved in lipid metabolism, and for cell wall biosynthesis (MMAR_1778: *tesA*), survival during infection (MMAR_0713: *pknG*) and the *esx-1* operon, as observed in *D. discoideum*, meaning that these genes are important for the survival of the bacteria in both hosts. Some mutations were specific to the BV2 microglial host, such as MMAR_0644, MMAR_1138, MMAR_2991, MMAR_3347, MMAR_4937, but their functions are not described except for a glucose-6-phosphate isomerase (MMAR_4557: *pgi*) and a trehalose-6-phosophate synthetase (MMAR_4978: *otsA*). However, in contrast to 96 hpi in *D. discoideum*, we did not find any mutation in genes belonging neither to the *mce1* nor to the *mce4* operon (Table S4).

## CONCLUSIONS

The recent combination of transposon insertion mutagenesis with “deep sequencing” renders it possible to quantify the abundance of insertion mutants in a pool to identify essential genes and genes required for selectable phenotypes (van Opijnen & Camilli, 2013). In the present study, we used the Tn-Seq approach to determine the fitness cost of mutations in *M. marinum* to obtain a comprehensive overview of host-pathogen interactions in two evolutionary distinct hosts, *D. discoideum* and BV2 microglial cells.

Our basic findings show that 57% of TA sites in the *M. marinum* genome were disrupted among which 9.2% of non-permissive insertion sites were identified. These results are in the same range as previous Tn-Seq studies on other pathogenic mycobacteria where around 50% of the TA sites were disrupted (Tateishi et al., 2020; J. Wang et al., 2014; Y. J. Zhang et al., 2012). More precisely, this result is comparable with a similar Tn-Seq study performed using *M. tuberculosis* which showed a range of 42% to 64% of TA sites mapped and 9% of non-permissive sites (DeJesus, Nambi, et al., 2017). The Mariner *Himar1* transposon used in this study has an absolute requirement for TA bases, therefore the high percentage of GC content in Mycobacteria genomes limits the potential of insertion, leading to a weaker coverage compared to other bacteria (Chao et al., 2013; Gawronski et al., 2009). Our study shows that 568 genes (10.2%) are essential for *M. marinum*, which is comparable to previous Tn-Seq studies on *M. tuberculosis*, in which 11.6% of the genes were essential (DeJesus, Gerrick, et al., 2017; Griffin et al., 2011). Surprisingly, another Tn-Seq conducted on the *M. marinum* E11 strain showed only 6% of essential genes (Weerdenburg et al., 2015). The difference between the two studies is probably due to their genomic diversity, a recent comparative genomic analysis revealed that *M*. *marinum* strains should be divided into two different clusters, the “M”- and the “Aronson”-type (Das et al., 2018). Among the essential genes identified in *M. marinum* M, 237 orthologues were found in *M. marinum* E11 (37.3%) (Weerdenburg et al., 2015), and 311 orthologues in *M. tuberculosis* (44.4%) (DeJesus, Gerrick, et al., 2017). The larger overlap with *M. tuberculosis* H37Rv compared to the E11 strain of the same species might be partly due *M. marinum* M strain and *M. tuberculosis* H37Rv being able to infect human, whereas *M. marinum* strain E11 has more environmental hosts. We identified 153 (2.2%) essential genes that were specific to the *M. marinum* M strain compared to *M. tuberculosis* or *M. marinum* E11. A possible explanation for this particularity might reside in the sequencing depth of the various studies or by the species-specific differences between the Mycobacteria genomes. Indeed, *M. marinum* M has a larger genome than *M. tuberculosis*, 6 Mb compared to 4 Mb (Stinear et al., 2008). In addition, our sample preparation protocol includes a PCR step with 25 cycles compared to 22 cycles in the other studies. We conclude without a big surprise that the “essential core” of *M. marinum* genes, is highly conserved with other pathogenic mycobacteria. Tn-Seq is a robust and statistically relevant technique to identify a range of pathways and mutations that have an impact on the bacterial fitness during infection. However, we cannot exclude that for some mutants the “helper effect” can occur partly due to the “clumpiness” of mycobacteria. As a result, a few bacteria can co-infect the same host cell during spinoculation and might even reside in the same MCV. Therefore, mutants can possibly rescue each other’s mutation, or aggravate them by negative epistatic interaction which might result in false positive or negative hits in terms of impact on the fitness cost.

The major pathways involved in the survival of *M. marinum* as highlighted by Tn-Seq during infection of *D. discoideum* are related to vitamin metabolism. Our findings suggest an important role of biotin (or vitamin B8) for *M. marinum* survival, with a strong FD when mutations occurred for example in the *MutA/B* and *BioA/B/F1/D* genes. Indeed, biotin is an essential cofactor for biotin-dependent enzymes involved in metabolic pathways such as the catabolism of fatty acids mediated by MutA/B in the TCA cycle (Salaemae et al., 2011). Interestingly, *M. tuberculosis* cannot scavenge sufficient biotin from its host, instead it uses *de novo* synthesis via genes such as *BioA, BioD*. Mutations in genes belonging to the biotin synthesis pathway which lead to FD, is in accordance with previous studies showing that biotin plays a role in the establishment and persistence of chronic infections in a murine model of tuberculosis (Woong Park et al., 2011). In addition, the biotin synthesis pathway is considered an interesting target to develop antibacterial agents, since mammals are not able to synthesise biotin, thus limiting the effects of synthesis inhibitors on the host (Bockman et al., 2020). Other important pathways for the survival of *M. marinum* are the carbon metabolism including glycolysis and gluconeogenesis. Mutations in genes belonging to these pathways had a fitness cost during infection, including *pckA*, encoding the first enzyme in gluconeogenesis, which is essential to establish and maintain infection (Marrero et al., 2010). On the other hand, our data highlight the importance of the methylcitrate cycle in establishment of Mycobateria infection, because mutations in *icl* lead to FD also during infection in *D. discoideum* (Muñoz-Elías & McKinney, 2005). Finally, one of the most studied and well conserved virulence pathways in Mycobacteria, the type VII secretion system ESX-1 encoded by the *esx-1* operon was identified in this study. Secretion of the *M. marinum* and the *M. tuberculosis* ESAT-6 and CFP-10 dimer, leads to MCV damage and finally bacterial escape to the cytosol (Conrad et al., 2017; van der Wel et al., 2007). All together, these findings validate that the Tn-seq method in *M. marinum* is a robust tool to highlight important pathways involved in the survival of the bacterium during infection. These results also demonstrate that the virulence strategies used by *M. marinum* to infect amoebae and mammalian cells are well conserved.

However, a few differences between the *D. discoideum* and BV2 models were observed. For instance, the heparin-binding hemagglutinin, HBHA initially described to play a role in adherence to non-phagocytic cells (Menozzi et al., 1996) and recently described to be involved in the formation of cytosolic lipid inclusion (Raze et al., 2018) was detected only in *D. discoideum*. In a similar way, genes encoding enzymes involved in mycobactin production, a siderophore produced by the bacteria to scavenge iron, was also found only in *D. discoideum* (Knobloch et al., 2020; Quadri et al., 1998). On the other hand, some pathways have been identified only in BV2 microglial cells model such as the PhoY2/PhoT two-component system involved in response to environmental stress including antibiotic treatment (Namugenyi et al., 2017; C. Wang et al., 2013) or the TrcR/TrcS two-component system expressed during growth within animal phagocytes (Haydel et al., 1999; Haydel & Clark-Curtiss, 2006).

These variations might be explained by differences in the time of selection, because only one timepoint at 48 hpi was used for BV2 microglial cells model compared to 96 hpi for *D. discoideum*. It is also important to note that the BV2 infections are performed at 32 °C compared to 25 °C for the *D. discoideum* infections. Finally, the difference might also be explained from a biological perspective, because different host cells use different strategies to combat infection, forcing *M. marinum* to respond/adapt by using slightly different virulence strategies.

Our Tn-Seq data show that mutations occurring in many sites of the *mce1* operon in *D. discoideum* model led to a FD whereas mutations in the same operon had a neutral effect during infection of BV2 microglial cells. In addition, mutations in genes encoding proteins involved in the stability of both the Mce1 and the Mce4 complex, such as the accessory OmamA protein (Perkowski et al., 2016) and the ATP-ase MceG, lead to a FD in *D. discoideum* only, while mutations in LucA had neutral effect in both phagocytic models (Nazarova et al., 2017). These results support the hypothesis addressed in the literature that mutations in the *mce1* operon impact fatty acid acquisition and lead to a decreased intracellular growth of *M. tuberculosis*, although a direct link between lipid acquisition and intracellular growth is not completely deciphered. One hypothesis of fitness cost phenotype variation between both phagocytes might be the difference in lipid composition between *D. discoideum* and BV2 microglial cells. *M. marinum* has the capacity to infect a wide range of host and therefore adapt its lipid metabolism according to lipids available in the infected milieu. To address this hypothesis, further investigations are required to understand the impact of *mce1* mutation during the infection course.

To conclude, this study reports a successful Tn-Seq performed in *M. marinum* M as a robust method to investigate fitness cost during infection. We identified several genes that potentially contribute to *M. marinum* infectivity that can be explored in future investigations. Taken together, this study opens the door for Tn-seq to be used to rapidly screen and identify genes involved in *M. marinum* fitness under a large variety of both *in vivo* and *in vitro* conditions.

## MATERIALS AND METHODS

### Bacterial strains, plasmids and culture methods

*M. marinum* M strain was grown in Middlebrook 7H9 (Difco) supplemented with 10% OADC (Becton Dickinson), 0.2% glycerol (Biosciences) and 0.05% Tween 80 (Sigma) at 32°C in shaking culture at 150 rpm in the presence of 5 mm glass beads to prevent aggregation. The axenic *D. discoideum* Ax2(Ka) was cultured at 22°C in Hl5c medium (Formedium) supplemented with 100 U/mL of penicillin and 100 μg/mL of streptomycin (Invitrogen). BV2 cells were grown at 37°C in DMEM High Glucose (Dulbecco’s Modified Eagle’s Medium) supplemented with 6**%** of heated inactivated Fetal Bovine Serum (FBS) and 100 U/mL of penicillin in 5% CO_2_ environment.

### Selection of transposon *M. marinum* mutants in *D. discoideum*

A transposon mutant library of *M. marinum* strain M was kindly obtained from Dr Graham Stewart’s laboratory. Mutants were generated as described previously (Butler et al., 2020; Long et al., 2015). The first round of selection was performed using aliquots of 6.5×10^4^ *M. marinum* transposon mutants expressing mCherry. The *M. marinum*-Tn library was thawed and grown overnight until an optical density (OD) of 0.8 was reached, centrifuged and resuspended in Hl5c. The inoculum was plated on ten 15-cm 7H10 plates supplemented with 25 μg/mL kanamycin, 10% OADC, 0.2% glycerol and 0.05% Tween-80. Following 7 days of incubation, the library was scraped from plates and identified as input samples for each experiment and stored in 10% glycerol at −80°C. For library selection, the *M. marinum*-Tn library inoculum was spinoculated on 5 x 10^6^ adherent *D. discoideum* cells at a multiplicity of infection (MOI) of 5 to avoid any helper effect by having 1 to 3 bacteria per cell. After washing off non-phagocytosed bacteria, the infected cells were incubated at 25°C in filtered Hl5c medium containing 5 μg/mL of streptomycin and 5 U/mL of penicillin to avoid extracellular growth. At either 6, 24 or 48 hpi *D. discoideum* were scraped and lysed with Triton 0.1%, intracellular bacteria were collected and plated on ten 15-cm 7H10 plates with 25 μg/mL kanamycin, 10% OADC, 0.2% glycerol and 0.05% Tween-80. The same incubation time was applied after each collected timepoint. The same procedure was used for the second round of selection but the inoculum used for this step was the frozen aliquot of *M. marinum* collected at 48 hpi after the first round.

### Construction of gDNA libraries of *M. marinum* mutant pools

*M. marinum* gDNA was extracted following an adapted protocol from Belisle et Sonnenberg (Belisle 6 Sonnenberg, 1998). Briefly, *M. marinum* mutant pools were centrifuged and resuspended in breaking buffer (50mM Tris, 10mM EDTA, 100mM NaCl, pH8). Mechanical lysis was performed with the Fastprep homogenizer during 30s, this step was repeated 3 times. The upper phase was chemically lysed with RNase (200μg/ml) and Lysozyme (100μg/ml) 1h at 37°C. Then, 0.1 volume of SDS 10% and 0.01 volume of Proteinase K (100μg/ml) were added for 1h at 55°C. Extraction steps were realized with equal volume of phenol:chlorophorm:isopropanol 25:24:1 for 30 min followed by chlorophorm:isopropanol during 5 min. Precipitation was performed with 3M sodium acetate pH 5.2 and 1 volume of isopropanol 100 %, gDNA was pelleted and washed with ethanol 70%. The extracted gDNA was quantified using Qubit (Invitrogen) and its quality was checked with the TapeStation system (Agilent).

A 2μg aliquot of gDNA in Tris Buffer (10mM Tris, 0.1mM EDTA, pH8) was sheared in a Covaris S2 machine following these parameters: Intensity 4, Duty cycle 10%, cycles 200 and time 65s. After purification with AMPure XP purification kit (Agencourt), 1000 ng of sheared DNA fragment ends were repaired, blunt ended and a d-A Tail added following NEBNextEnd/d A-Tailing module protocol. Specific adapters 1 and 2 described in Table B were self-hybridized and then ligated to the ‘A’ tail ends by T4 ligase (Promega). After purification with AMPure XP purification kit (Agencourt), 40 ng of libraries were amplified by PCR using KAPA polymerase with the following parameters: 95°C 3 min; 98°C 20s, 60°C 15s, 72°C 15s (repeated for 25 cycles); 72°C 1 min for elongation. Transposon junctions were amplified with the IS6 primer: 5’ CAAGCAGAAGACGGCATACGA 3’ and specific primers MarA to MarP described in Table B. PCR products were purified with AMPure XP purification kit and Sixteen-plexed samples were pooled in approximately equimolar amounts of 5 nM and run in a single and double index reads on a Hiseq 4000 (Illumina).

### Transposon Sequence Analysis

The first step includes the pre-processing of the obtained raw sequences. The raw sequence files are firstly checked using the FASTQC tool which gives the overall sequence quality plots by which the low-quality samples can be detected and outliers can be discarded if necessary. For each read, the read end nucleotides with FASTQ quality Phred score will be chopped, and the whole read will be disposed if number of ‘#’ > 50bp of half read length. Also, nontransposon reads are filtered out by keeping the reads containing the sequence ‘TGTTA’ at the beginning of the read. Following the pre-processing step, reads were mapped on the reference genome, which consists of the sequence of *M. marinum* M genome and *M. marinum* M plasmid pMM23 using Bowtie2. Only alignments with mapping quality > 30 are kept for the downstream analysis which consists of counting transposon insertions from alignment SAM files. Alignments having the same alignment location and indexed code were considered as PCR duplicated alignments and subsequently removed prior to counting of transposon insertions for each TA site. The gene essentiality analysis was conducted using the TRANSIT tool version which includes HMM and re-sampling methods (DeJesus et al., 2015). Briefly, the HMM method predicts essentially based on the entire genome in a single condition, whereas the resampling method is a comparative analysis which can be used to determine conditional essentiality of genes between two conditions. With TRANSIT, the essentiality of genes was evaluated separately for each condition. The essential core was determined by taking the intersection of genes with an essential call “ES” over all tested conditions. One gene was removed, due to a lacking MMAR ID. The essential core from this study was compared with genes being identified as essential in two other studies (DeJesus, Gerrick, et al., 2017; Weerdenburg et al., 2015). Only orthologous genes were considered (E11 orthologues given by Weerdenburg et al. (Weerdenburg et al., 2015), H37Rv orthologues given by mycobrowser.epfl.ch, Release 3, 2018-06-05) and only *M. marinum* M strain genes with a MMAR ID were used. This was done using R 4.2.1.

As a result, significant genes can be selected according to fold change values and *p*-values from re-sampling results. R package limma was used to perform the differential expression analysis on insertion count per gene. The raw count data are firstly normalized to obtain the same total counts across samples, then the read counts were log-transformed using voom into log2-counts per million (log_2_CPM). Apart of tools described above, all analyses were performed using own developed scripts.

### Cluster analysis for Evolution of the GD/GA scores

The gene clusters analysis was performed with the R programming language, using the content of Table S2 as input. Genes with at least 5 TA sites and a normalized read count > 1 in any of the samples were selected. For each gene, we compute its profile across timepoints by considering the average of the normalized count for each condition. The Manhattan distance between each pair of gene-profile was computed and a hierarchical clustering algorithm with ward.D linkage was run. The resulting clustering tree was cut at a height of 35, allowing to retain 9 clusters (Table S3).

### Gene Ontology analysis

Before functional group analysis via KEGG and GO enrichment, genes were thresholded as follows: Fitness advantage (FA), log_2_fc ≥ 0.585, Fitness disadvantage (FD), log_2_fc ≤ −0.585, FA and FD, abs(log_2_fc) ≥ 0.585. Genes were additionally thresholded by *p*-value as follows: 24 hpi and 48 hpi, *p*-value ≤ 0.1, 96 hpi, *p*-value ≤ 0.01. Different significance thresholding was motivated by better visualization of enriched terms.

Prior to submission to the enricher function or enrichKEGG function of the R package clusterProfiler, only genes with a MMAR ID were used and mapped to UniProt AC using the mycobrowser annotation (Release 3, 2018-03-28) and the UniProt annotation (queried for “Mycobacterium marinum”, https://doi.org/10.1093/nar/gkaa1100).

GO and KEGG term analysis was done with R (version 4.2.1) and the package clusterProfiler (version 4.4.4, 10.1016/j.xinn.2021.100141). For KEGG analysis the KEGG annotation (Release 103.1, September 1, 2022, https://doi.org/10.1093/nar/28.1.27) version was used, for GO terms, the QuickGO annotation for *Mycobacterium marinum* (version from 19.03.2021, 10.1093/bioinformatics/btp536) was used. GO term names were extracted using the R package GO.db version 3.15.0. Group size was limited between 2-200 genes and only filtered by *p*-value < 0.01 for GO term analysis and *p*-value < 0.05 for KEGG analysis. For GO term analysis, all three aspects of the ontology were used, biological processes, molecular function, and cellular component.

### STRING database analysis

The analysis was performed using Cytoscape 3.9.1 (Shannon et al., 2003) with Omics Visualizer 1.3.0 (Legeay et al., 2020) and the STRING database version 11.5 (Szklarczyk et al., 2015). First, log_2_fc and *p*-values were taken from comparing 24, 48 and 96 hpi to the inoculum. Genes were filtered for absolute log_2_fc ≥ 0.585, *p*-value ≤ 0.05 in any of the three timepoints and converted to each respective ortholog in *Mycobacterium tuberculosis*, for which the full STRING network was subsequently retrieved. The network was cut at a confidence cut-off of 0.8 and singletons were omitted. The colour scale in the layered rings represents from inside to outside the log_2_fc of 24, 48 and 96 hpi respectively compared to the inoculum.

## Abbreviation list

Mm: *Mycobacterium marinum*,
Mm-M: *Mycobacterium marinum* Strain M,
Mm-E11: *Mycobacterium marinum* Strain E11,
Mtb: *Mycobacterium tuberculosis*,
Dd: *Dictyostelium discoideum*,
BV-2: murine microglial cells,
GA: Growth advantage,
GD: Growth disadvantage,
FA: Fitness advantage,
FD: Fitness disadvantage,
NE: Neutral,
ES: Essential

## ACKNOWLEDGEMENTS

We acknowledge Dr. Huihai Wu for his help in analysing the the TnSeq data, the staff from the core facilities IGE3 Genomics platform at the Faculty of Medicine for their precious help. We thank Dr H. Koliwer-Brandl and Cristina Boehm-Bosmani for their involvement in various preliminary experiments. This work was supported by multiples grant from; the National Centre for the Replacement, Refinement and Reduction of Animals in Research (NC3Rs) (awarded to GRS and TS), the SystemsX.ch grant HostPathX to TS, and the Swiss National Science Foundation research grants 310030_169386 and 310030_188813 to TS.

---

**Table S1 Insertion counts per TA site for *D. discoideum* and BV-2 experiments**

**Table S2 Gene counts for *D. discoideum* and BV-2 experiments**

**Table S3 Conditional essentiality analysis for *D. discoideum* and BV-2 experiments**

**Table S4 Enrichment analyses for *D. discoideum* and BV-2 experiments**

## Supplementary Figures

**Fig S1. PCA of all experimental conditions in *D. discoideum***

Principal components 1 and 2 of normalized insertion counts, colored by conditions: Inoculum (n=7), 6 hpi (n=4), 24 hpi (n=6), 48 hpi (n=8), Inoculum Bis (n=6) and 96 hpi (n=6). Below the axis label the percentage of variance captured by the respective principal component is denoted.

**Fig S2. Differential representation of *M. marinum* mutants**

**A.** Essential genes for each condition were determined with TRANSIT. The Venn diagram shows the intersection of all four conditions, i. e. the “essential core” containing 568 genes.

**B.** KEGG terms enriched in the essential core sorted as a dotplot. Using R and the package clusterProfiler terms were filtered by *p*-value < 0.05. The *p*-value is color coded, and the number of enriched genes found in the respective term are coded by dot size. Edges between nodes represent overlap between the connected terms.

**Fig S3 Hierarchical clustering of log2 fold changes in *D. discoideum***

**A, B, C**. Enriched GO terms in cluster 2, 7 and 9 depicted as dot plot. Using R and the package clusterProfiler terms were filtered by *p*-value < 0.01. The *p*-value is color coded, and the number of enriched genes found in the respective term are coded by dot size.

**Fig S4. Fitness advantage (FA) and fitness disadvantage (FD) over the infection time course in *D. discoideum***

Genes thresholded as described in Table 1 were submitted to GO term enrichment using R and the package clusterProfiler. Terms were filtered by *p*-value < 0.01 and visualized as dotplots. The *p*-value is color coded and the number of enriched genes per term is coded by the dot size.

**Table A.**
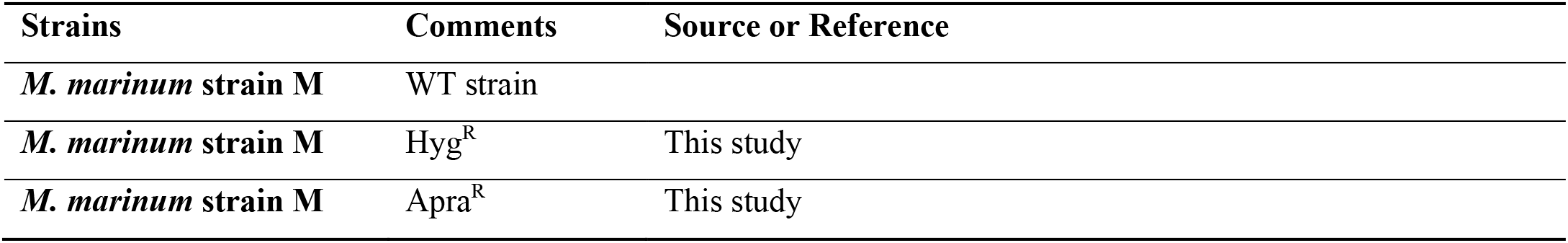
Bacterial strains used in the study.

| Strains | Comments | Source or Reference |
| --- | --- | --- |
| *M. marinum* strain M | WT strain |  |
| *M. marinum* strain M | Hyg<sup>R</sup> | This study |
| *M. marinum* strain M | Apra<sup>R</sup> | This study |

**Table B.**
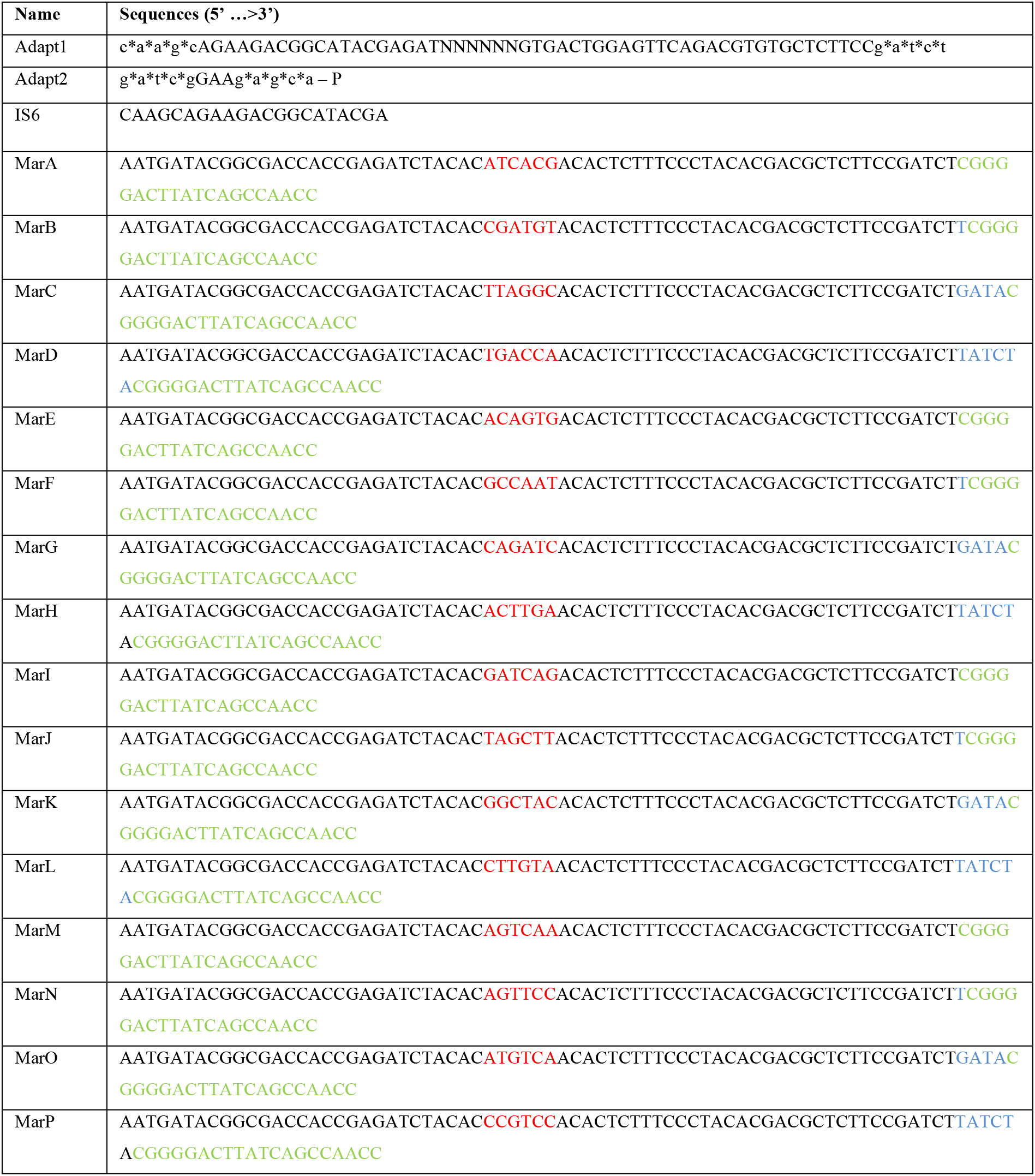
Primers used in the study for *M. marinum* mutant pools amplification.

**A.** Enriched GO terms for FA and FD at 24 hpi **B.** Enriched GO terms FA and FD at 48 hpi **C.** Enriched GO terms FA at 96 hpi **D.** Enriched GO terms FD at 96 hpi. **E, F, G** show volcano plots of binary comparisons of 24 hpi, 48 hpi and 96 hpi respectively to the Inoculum. The x-axis shows the log_2_fc and the y-axis the -log^10^ of the associated *p*-values. The lowest thresholds used to filter genes for enrichment analysis are depicted as thick black lines: log_2_fc ≥ 0.585 and ≤ 0.585 (corresponding to 1.5 fold change) and *p*-value ≤ 0.1. For visualization *p*-values < 0.0001 were capped to 0.0001.

**Fig S5. Clustering of differentially affected genes of *M. marinum* during infection of BV2 microglial cells**

**A.** Plots of principal component analysis of the *M. marinum* Inoculum (n=4) versus 48 hpi (n=3) of BV2 infection. **B.** For all genes with at least 10 TA sites, we have considered the proportion of TA inserted with more than 1 TA with normalized read count. The distribution for each sample are reported and grouped by different colors according to the 2 conditions used for the analysis (Inoculum versus 48 hpi).

