## Supplementary Figures for "Temporal genome-wide fitness analysis of *Mycobacterium marinum* during infection reveals genetic requirement for virulence and survival in amoebae and microglial cells"

Lefrançois et al., Figure S1

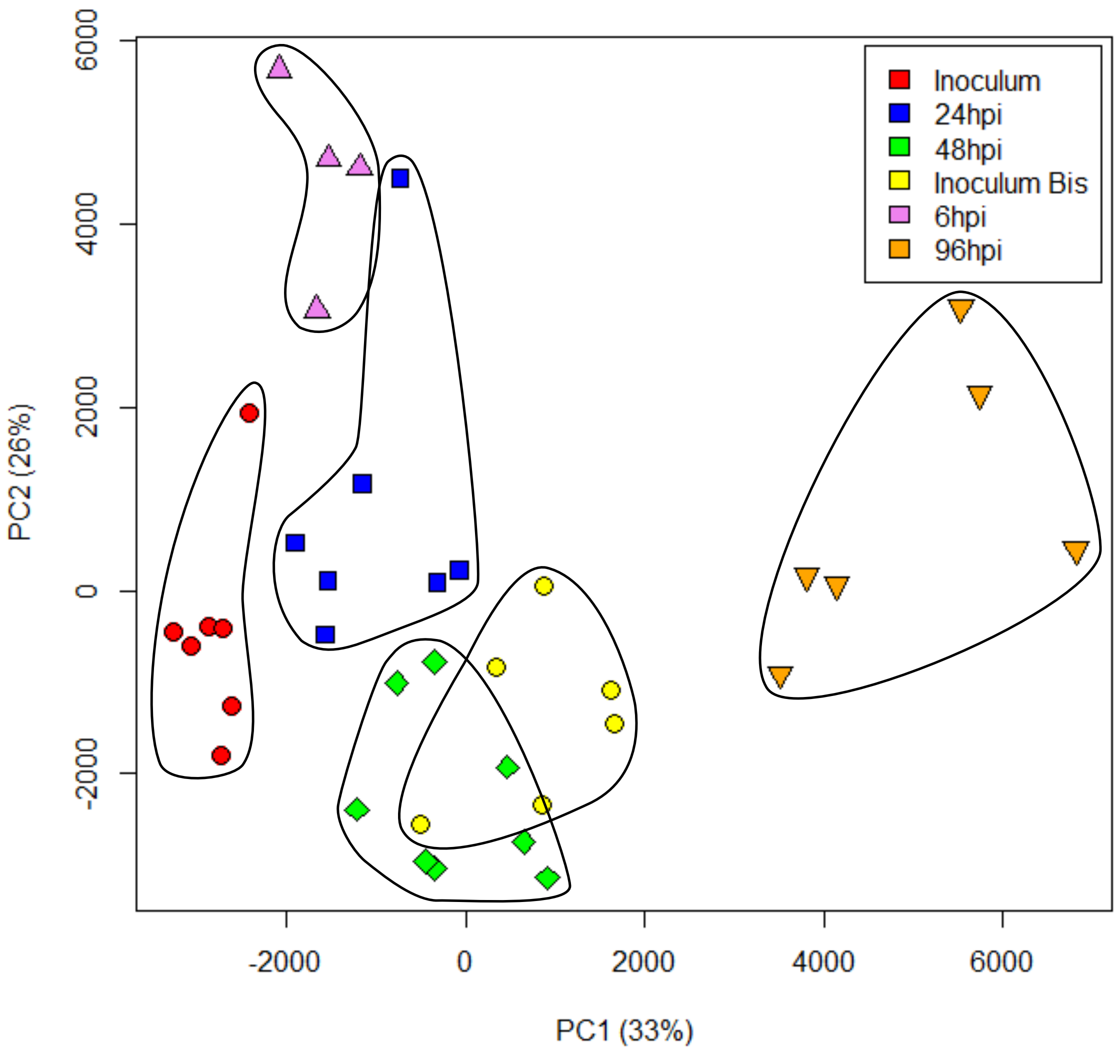

Lefrançois et al., Figure S2

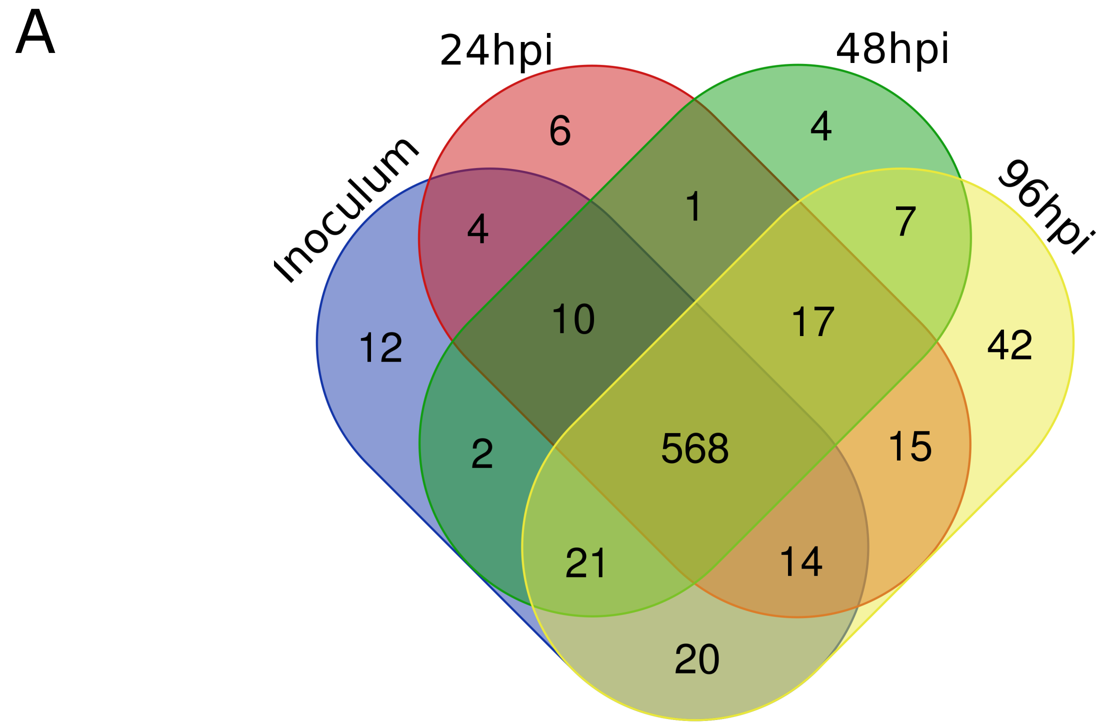


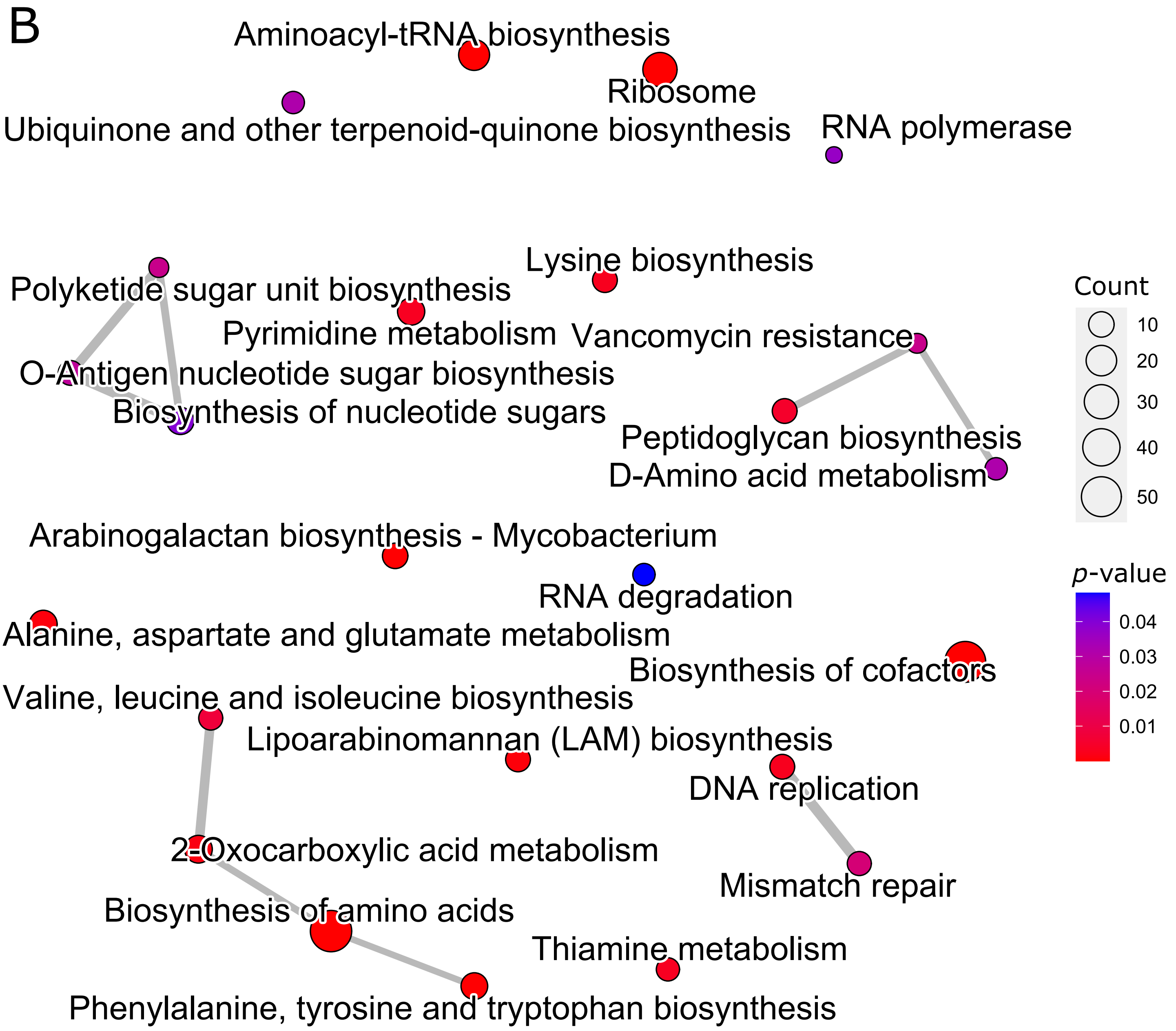

Lefrançois et al., Figure S3

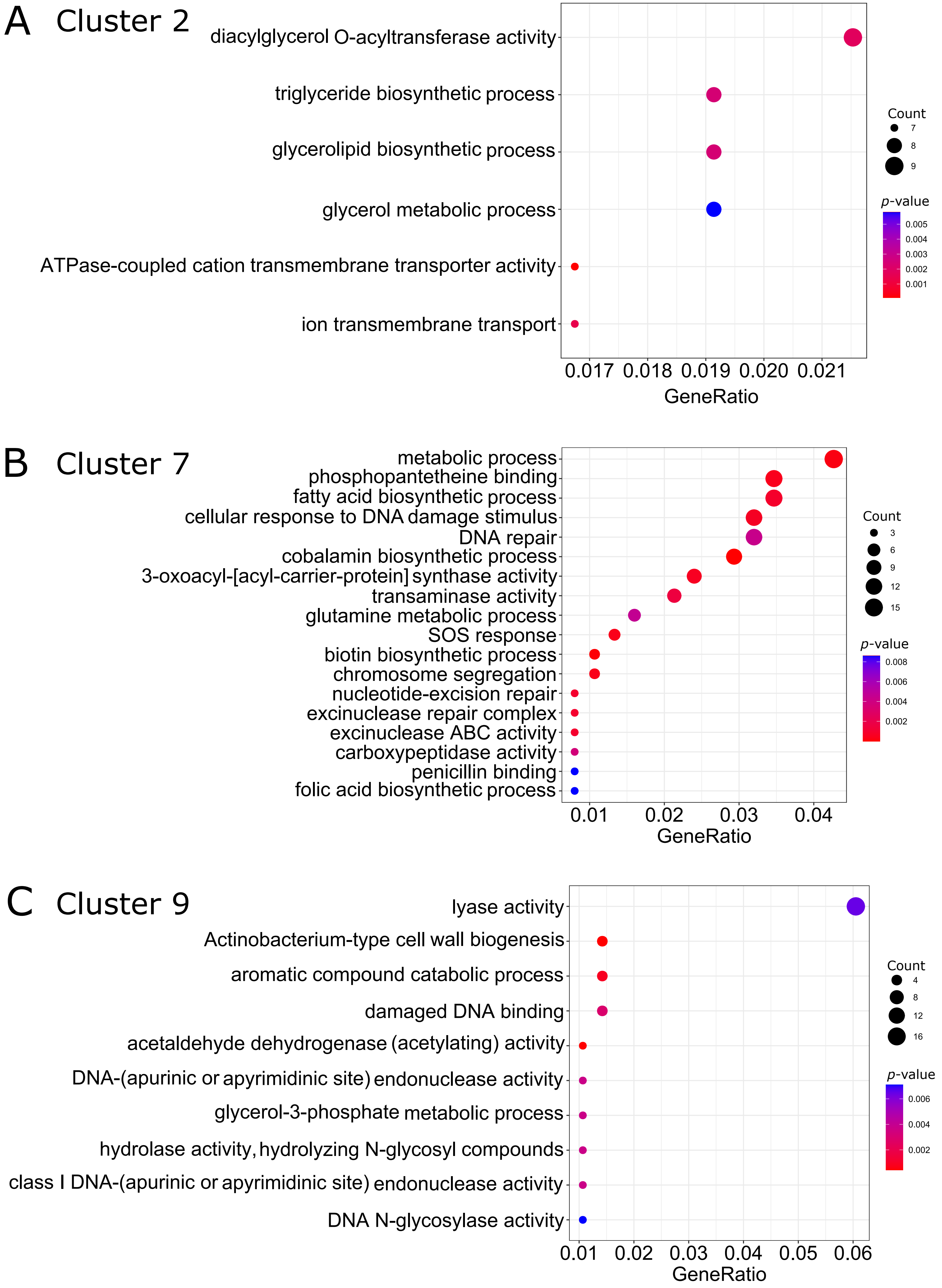

Lefrançois et al., Figure S4

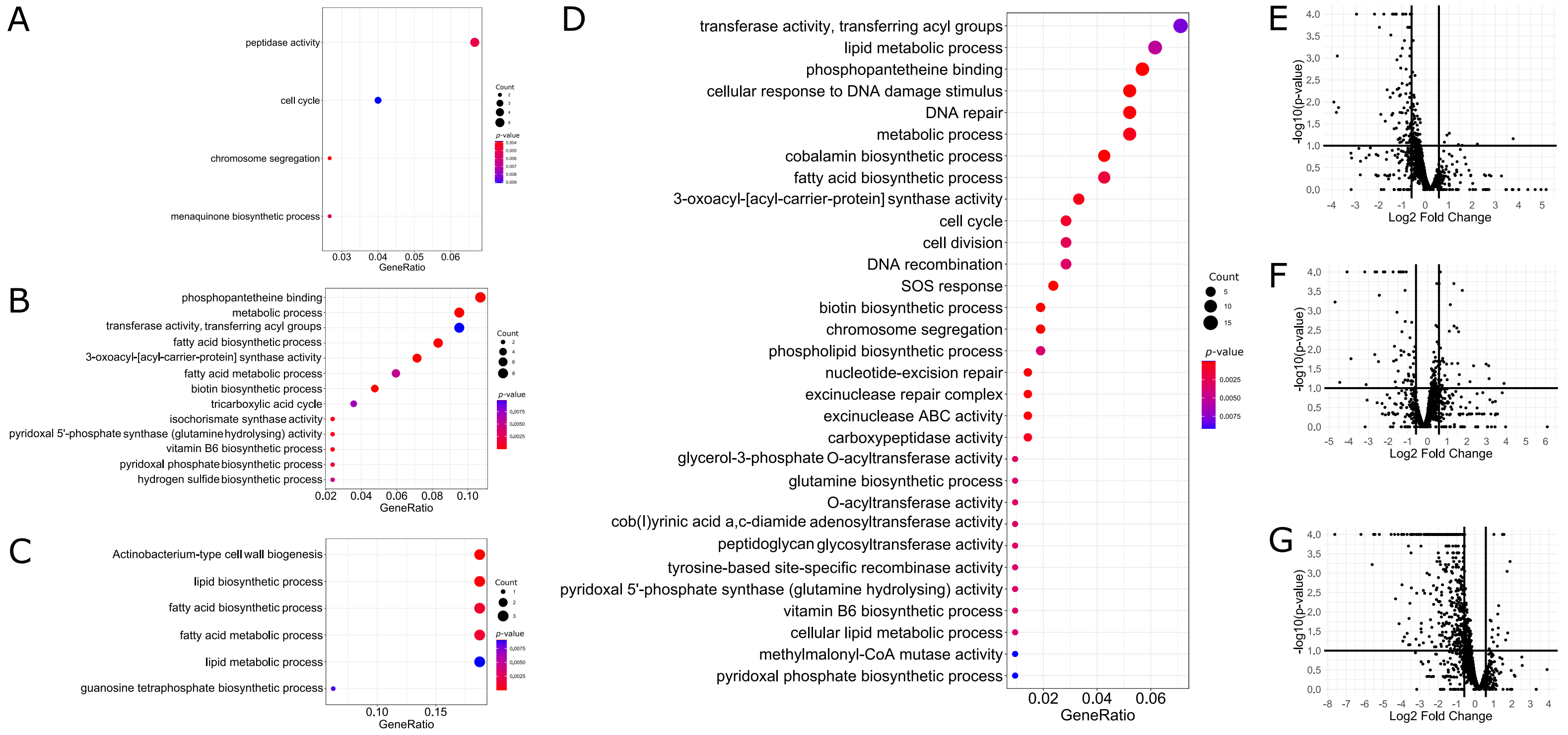


H

| | | FA and FD | FA | FD |
|---|---|---|---|---|
| *D. discoideum* | 24 hpi | 97 | 7 | 90 |
| | 48 hpi | 97 | 38 | 59 |
| | 96 hpi | 268 | 19 | 261 |
| BV-2 | 48 hpi | 183 | 12 | 171 |

Table 1: Differentially transposon mutated genes after different periods of infection. Gene log2 fold changes (log2fc) with respect to the inoculum were thresholded as follows: Fitness advantage (FA), log2fc $\geq$ 0.585, Fitness disadvantage (FD), log2fc $\leq$ -0.585, FA and FD, abs(log2fc) $\geq$ 0.585. For data obtained from infected *D. discoideum* genes were additionally thresholded by *p*-value as follows: 24 hpi, 48 hpi and FA at 96 hpi *p*-value $\leq$ 0.1, 96 hpi, p-value $\leq$ 0.01. For data obtained infected from BV-2 microglial cells genes were thresholded by *p*-value $\leq$ 0.1. Different significance thresholding was motivated by better visualization of enriched functional groups.

Lefrançois et al., Figure S5

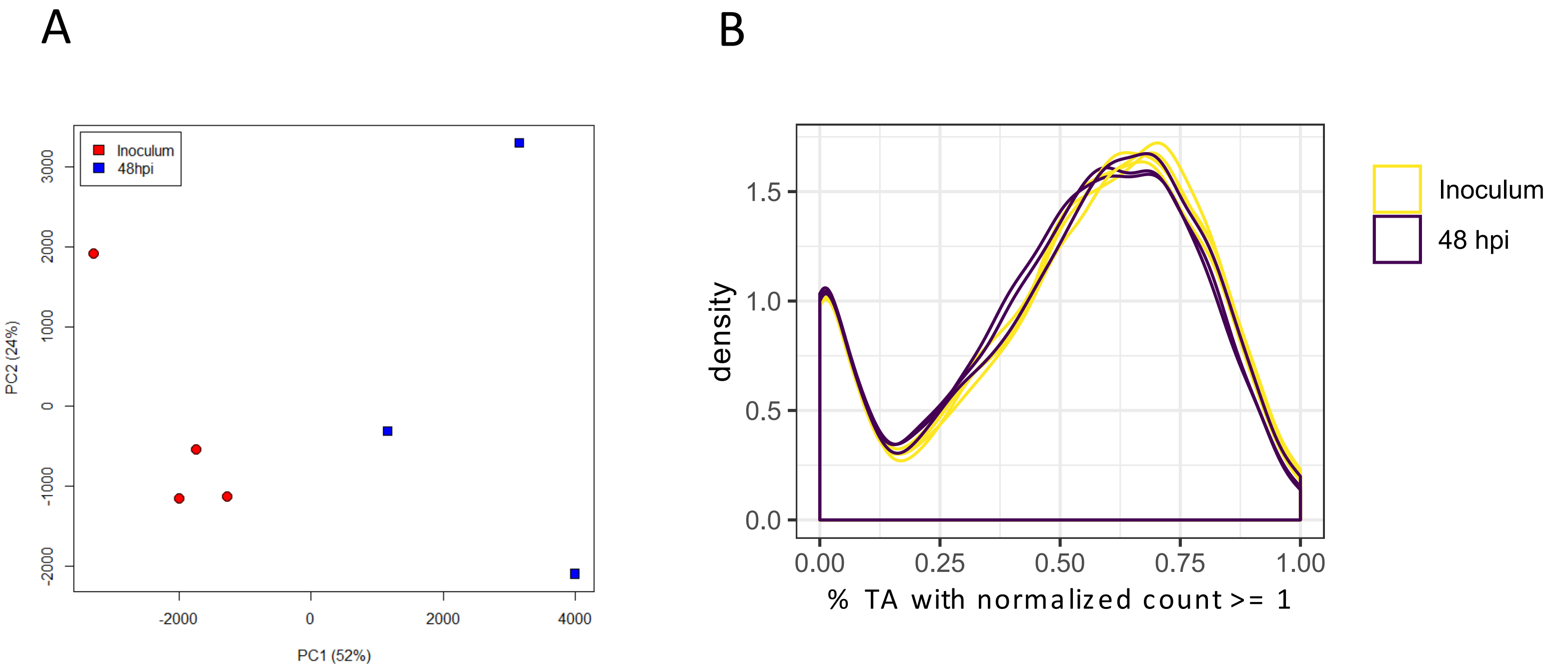
